# ARHGEF17 Deficiency Induces Endothelial Dysfunction and Intracranial Aneurysm Formation via RhoA/ROCK2/MLC Signaling Pathway

**DOI:** 10.1101/2025.10.14.682476

**Authors:** Jian Li, Hao Zhang, Chao Peng, Bangyue Wang, Yan Zhao, Xinyu Yang

**Affiliations:** Department of Neurosurgery, Tianjin Medical University General Hospital; Neuroendocrinology Laboratory Tianjin Institute of Neurology; Nanjing Drum Tower Hospital, The Affiliated Hospital of Nanjing University Medical School; Southwest Hospital, Army Medical University, Chongqing, China; The Second Affiliated Hospital of Anhui Medical University

## Abstract

**BACKGROUND:** Genetic susceptibility is a major determinant in intracranial aneurysm (IA) formation and rupture, yet the underlying mechanisms linking genetic variation to vascular dysfunction remain largely unknown. We have identified mutations in ARHGEF17, a guanine nucleotide exchange factor that regulates RhoA activation and cytoskeletal organization, as potential risk variants for IA. Given ARHGEF17’s regulatory role in endothelial barrier integrity and actin remodeling, we hypothesized that ARHGEF17 deficiency promotes IA pathogenesis through dysregulation of the RhoA/ROCK2/MLC signaling axis, leading to endothelial dysfunction and vascular wall instability.

**METHODS:** CRISPR–Cas9–mediated ARHGEF17 knockout (ARHGEF17^⁻/⁻^) mice and morpholino-based ARHGEF17-deficient zebrafish were established to assess the in vivo vascular effects of ARHGEF17 loss. An intracranial aneurysm model combining elastase injection and deoxycorticosterone acetate (DOCA)–induced hypertension was used to evaluate aneurysm incidence, rupture rate, and survival. Structural remodeling of the Circle of Willis (CoW) was assessed by Victoria Blue, EVG, and Picrosirius Red staining, as well as immunofluorescence for α-SMA, OPN, CD31, and inflammatory markers. Complementary in vitro studies were performed in HUVECs using lentiviral ARHGEF17 silencing (three shRNAs of varying efficiency) to examine endothelial proliferation, migration, tube formation, and barrier function (TEER). Activation of RhoA/ROCK2/MLC signaling was quantified by G-LISA and Western blotting. The ROCK inhibitor Y-27632 (10 μM) was applied to determine pathway dependence.

**RESULTS:** ARHGEF17^⁻/⁻^ mice exhibited a significantly higher incidence and rupture rate of intracranial aneurysms, accompanied by fragmentation of elastic fibers, loss of collagen organization, vascular smooth muscle cell dedifferentiation, and robust inflammatory activation in the CoW. Zebrafish lacking ARHGEF17 showed frequent intracranial hemorrhage and compromised vascular wall integrity, further confirming ARHGEF17’s role in cerebrovascular stability. In ECs, ARHGEF17 knockdown impaired proliferation, migration, tube formation, and barrier integrity in a silencing-efficiency–dependent manner. Mechanistically, ARHGEF17 deficiency activated the RhoA/ROCK2/MLC pathway, leading to increased phosphorylation of MLC and MYPT1 and disorganization of F-actin and junctional proteins. Pharmacological inhibition with Y-27632 restored endothelial function, normalized cytoskeletal structure, and re-established junctional continuity, indicating a ROCK2-dependent mechanism.

**CONCLUSIONS:** Our findings establish ARHGEF17 as a critical regulator of cerebrovascular integrity and identify RhoA/ROCK2/MLC mediated cytoskeletal remodeling as the mechanistic link between ARHGEF17 deficiency and aneurysm pathogenesis. Loss of ARHGEF17 compromises endothelial barrier function, triggers vascular inflammation, and promotes aneurysm formation and rupture. Importantly, ROCK inhibition rescues endothelial dysfunction, highlighting the RhoA/ROCK2/MLC axis as a promising therapeutic target for ARHGEF17 mutation–associated intracranial aneurysms.

## INTRODUCTION

Intracranial aneurysm (IA), which is characterized by the localized dilation of the cerebral artery, is a common vascular abnormality primarily affecting the Circle of Willis(CoW), with a prevalence of 2% to 5% in the general population and frequently leads to vascular rupture with high mortality^[1, 2]^. The pathogenesis of IA involves endothelial cells (ECs) dysfunction, vascular smooth muscle cell (VSMC) death, and extracellular matrix(ECM) disruption, which lead to aneurysm formation and rupture^[3]^. However, the aetiology and pathogenesis of IA are not fully understood.

Current treatment strategies primarily focus on endovascular interventions and surgical craniotomy, which are aimed at preventing IA progression. However, they do not address the underlying disease mechanisms and carry inherent risks and limitations, effective therapeutic strategies to alleviate disease deterioration and reverse disease progression are still lacking^[4]^. Therefore, elucidation of the molecular mechanism of this disease might identify optimal treatments.

A recent genome-wide association study(GWAS), which included the largest cohort to date, has identified Genetic predisposition is critical in IA and aneurysmal subarachnoid hemorrhage (aSAH)^[5]^. However, the mechanisms by which these single nucleotide polymorphisms (SNPs) affect the pathogenesis or progression of IAs remain to be elucidated^[5–7]^. Previously, based on the next-generation sequencing in a discovery cohort of Chinese IA patients, bioinformatics filters were exploited to search for candidate deleterious variants with rare and low allele frequency. We further examined the candidate variants in a multiethnic sample collection of 86 whole exome sequenced unsolved familial IA cases from 3 previously published studies. These findings indicate that ARHGEF17 mutations may be IA risk factors and have a functional basis in the integrity of intracerebral arteries^[8]^.

ARHGEF17, also known as tumor endothelial marker 4 (TEM4), is a member of the Dbl family of Rho guanine nucleotide exchange factors (RhoGEFs). It encodes the protein RhoGEF17. Along with GTPase accelerating factors, it regulates the GTP-bound state and, therefore, the activity of Rho GTPases. ^[9, 10]^. With GTP bound, Rho GTPase function as master switches controlling core cell functions, including tension and barrier function in endothelial cells^[8, 11, 12]^. In ECs, ARHGEF17 localizes to the actin cytoskeleton and cell–cell junctions via sequences in the N-terminus and activates RhoA at the intercellular junctions, where it plays a crucial role in maintaining barrier integrity and cytoskeletal organization, the depletion of ARHGEF17 leads to defective human umbilical vein endothelial cell junctions^[9, 13]^.

Moreover, it has been shown that the specific activation of RhoA at intercellular junctions is critical for proper endothelial junction formation and integrity^[14]^. Our studies on ARHGEF17 demonstrated that mutation of ARHGEF17 disrupts vascular endothelial integrity, enhances paracellular permeability, and leads to disorganized actin stress fibers, indicating its essential role in stabilizing intercellular junctions.

In this work, we highlight a critical role of the mutation of ARHGEF 17 in IAs and its potential clinical relevance as a prognostic and therapeutic biomarker in IAs. In our research, we demonstrate that ARHGEF17 alone is sufficient to drive the disruption of vascular integrity. By integrating functional and molecular studies in mouse, zebrafish and ECs, we show that ARHGEF17 leads to the upregulation of RhoA/Rock2 signaling to the actin cytoskeleton. This, in turn, results in changes in the actin cytoskeleton dynamics, leading to phosphorylation of MLC. These finding sheds light on the critical role of ARHGEF17 in perturbing the normal structure and function of the vascular system, highlight the RhoA/Rock2/MLC circuitry as a therapeutic vulnerability in ARHGEF17-mutation IAs patients that can be targeted pharmacologically to prevent progression to IAs.

## METHODS

Detailed methods are included in the online-only Data Supplement. The data that support the findings of this study are available from the corresponding author on reasonable request.

## Results

### ARHGEF17-Deficient induces IAs and promote rupture in IAs mouse models

To investigate whether deficient of ARHGEF17 increases the susceptibility of mice to IA formation, we constructed ARHGEF17 KO mice (ARHGEF 17^-/-^) to mimic the loss of ARHGEF 17 function using the CRISPR–Cas9 gene-editing system (Supplementary Fig. S1). ARHGEF17 knockout was validated and confirmed by agarose gel electrophoresis (Supplementary Fig. S2). Furthermore, we assessed the effect of ARHGEF17 deficiency on the formation and progression of intracranial aneurysms in an IAs mouse model^[15, 16]^.

Morphological examination of the CoW revealed that ARHGEF17^⁻/⁻^ mice pronounced vascular tortuosity and global arterial irregularities, with more extensive dilation and wall deformation throughout the CoW. In particular, the number and size of aneurysmal protrusions were markedly increased, and the regions of vascular rupture and subarachnoid hemorrhage exhibited clear expansion relative to WT controls (Fig.1 A-C). Moreover, the incidence of IA formation was significantly higher in the ARHGEF17^⁻/⁻^ group (76%) than in WT group (52%) after IA induction (n = 25, Fig.1D). Importantly, ARHGEF17 deficiency also increased the rate of aneurysms rupture (84.21% vs. 46.15%, Fig.1E). Kaplan–Meier survival analysis demonstrated a significantly lower survival rate in ARHGEF17^⁻/⁻^ IA mice compared with WT (log-rank test, p = 0.0034, Fig.1F). Quantitative morphometric analysis confirmed that the maximal luminal diameter of the CoW was significantly larger in ARHGEF17^⁻/⁻^ IA mice than in WT IA mice (230.5 ± 9.3 µm vs. 180.8 ± 8.6 µm, Fig.1G, H).

**Fig. 1.**
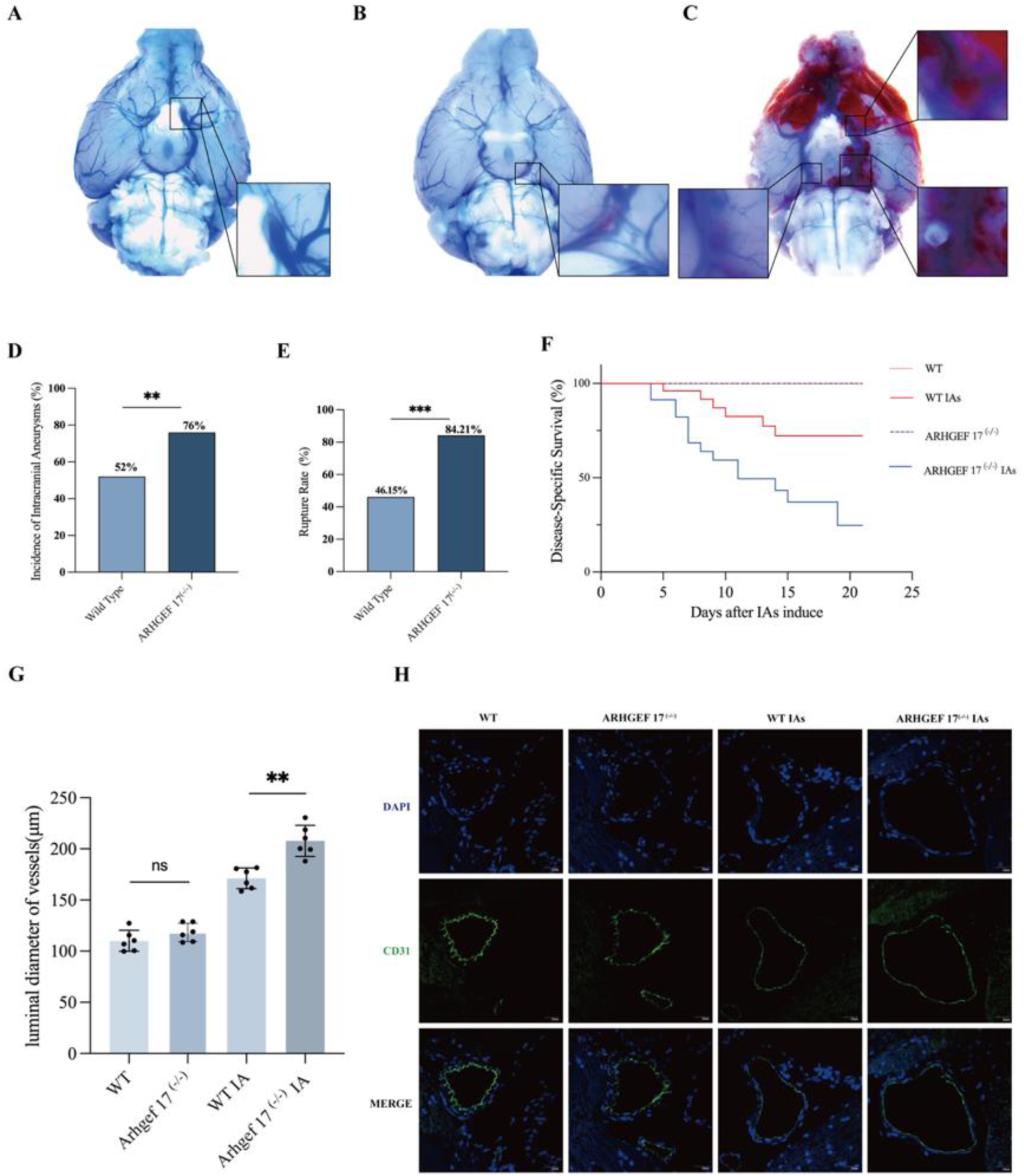
ARHGEF17 Deficiency Induces Intracranial Aneurysm Formation and Promotes Rupture in Mouse Models (A) Representative image of an unruptured intracranial aneurysm (IA) in a wild-type (WT) mouse, showing no clinical symptoms throughout the observation period. (B) A ruptured IA with subarachnoid hemorrhage observed in a WT mouse at 7 days post-induction. (C) Multiple ruptured and hemorrhagic aneurysms in an ARHGEF17^⁻/⁻^ IA model mouse, indicating more severe vascular injury and rupture. (D) Quantitative analysis of IA incidence in WT and ARHGEF17^⁻/⁻^ mice. The incidence was significantly higher in ARHGEF17^⁻/⁻^ mice (76%) than in WT controls (52%). (E) Comparison of IA rupture rates showing a markedly increased rupture incidence in ARHGEF17^⁻/⁻^ IA mice (84.21%) compared with WT IA mice (46.15%). (F) Kaplan–Meier survival analysis demonstrating a significant reduction in overall survival in ARHGEF17^⁻/⁻^ IA mice compared with WT IA mice during the follow-up period (log-rank test, P = 0.0034). (G) Quantitative measurement of the luminal diameter of the Circle of Willis (CoW) among groups. No significant difference was observed between normal WT and ARHGEF17^⁻/⁻^ mice (P > 0.05), whereas the diameter was significantly increased in ARHGEF17^⁻/⁻^ IA mice compared with WT IA mice (207.9 ± 8.6 µm vs. 171.2 ± 7.9 µm, P = 0.0045). (H) Representative immunofluorescence images of the CoW stained with CD31 showing vascular morphology and luminal dilation. Scale bars = 20 µm.

Together, these findings demonstrate that loss of ARHGEF17 profoundly enhances vascular vulnerability, leading to increased aneurysm formation, dilation, and rupture, ultimately resulting in poorer survival outcomes. To further elucidate the cellular and structural basis of this phenotype, we next examined the histopathological remodeling of the CoW in ARHGEF17^⁻/⁻^ mice.

### ARHGEF17-Deficient Triggers Abnormal Structural and Functional Remodeling of the CoW in Mice

Vascular structural deterioration is a hallmark of IA pathogenesis^[17]^.In the present study, ARHGEF17 deficiency markedly exacerbated this pathological process in the IA mice model. Histopathological staining analyses revealed profound vascular wall remodeling in ARHGEF17^⁻/⁻^ mice compared with WT counterparts with normal cerebral arteries (Fig.2). Victoria blue staining, which specifically visualizes elastic fibers, uncovered distinct structural differences between groups at the microscopic level. In WT mice, elastic fibers appeared as continuous, linear, blue-stained lamellae forming dense and parallel layers along the vascular wall, thereby maintaining vascular elasticity and integrity. In contrast, ARHGEF17^⁻/⁻^ IA mice exhibited extensive elastic fiber damage, characterized by marked discontinuity, focal fragmentation, and local absence of staining, leaving gaps within the vascular wall.

**Fig. 2.**
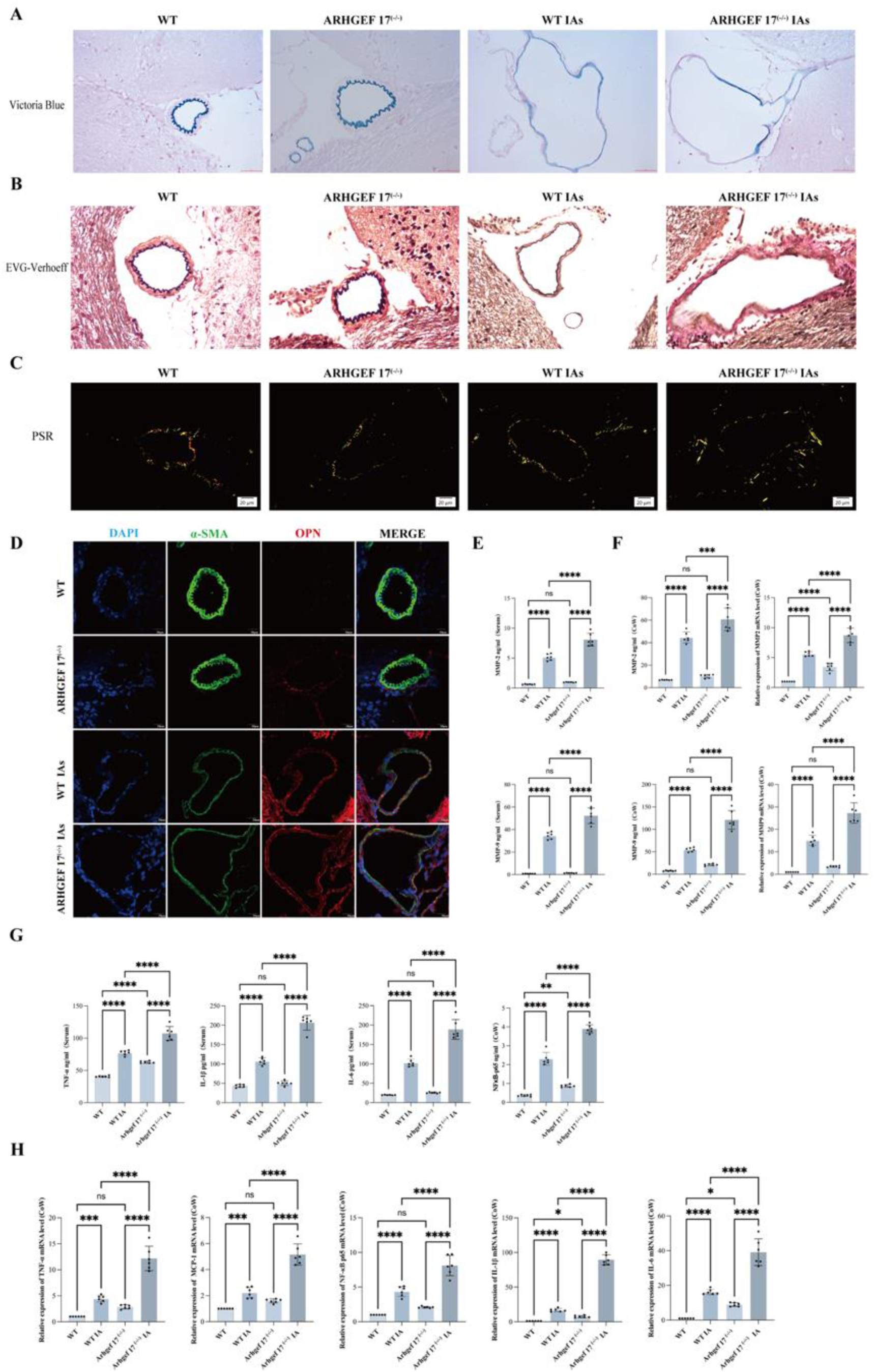
ARHGEF17 Deficiency Exacerbates Structural and Functional Remodeling of the Cerebral Arterial Wall in Intracranial Aneurysm (IA) Mice (A) Victoria Blue staining showing the organization of elastic fibers in the CoW. WT mice exhibited continuous, densely aligned elastic lamellae, whereas ARHGEF17^⁻/⁻^ IA mice displayed extensive elastic fiber fragmentation, discontinuity, and local loss of integrity, indicative of severe elastic fiber degradation. (B) Elastica van Gieson (EVG) staining demonstrating the architecture of elastic and collagen fibers. WT arteries exhibited a well-preserved internal elastic lamina (IEL) and compact collagen structure, while ARHGEF17^⁻/⁻^ IA arteries showed fragmented or absent IEL and loosely arranged, uneven collagen fibers. (C) Picrosirius Red (PSR) staining under polarized light revealing extracellular matrix remodeling. WT arteries showed dominant orange-red birefringence corresponding to mature type I collagen, whereas ARHGEF17^⁻/⁻^ IA arteries exhibited increased greenish-yellow birefringence indicative of immature type III collagen, reflecting collagen disorganization and ECM instability. (D) Immunofluorescence staining for á-smooth muscle actin (á-SMA) and osteopontin (OPN) showing VSMC phenotypic switching. WT and ARHGEF17^-/-^ arteries exhibited strong, continuous α-SMA signals along the tunica media, while ARHGEF17^⁻/⁻^ IA arteries displayed markedly reduced and discontinuous α-SMA staining vs WT IAs. OPN expression was significantly upregulated in the media and intima of ARHGEF17^⁻/⁻^ IA mice compared with WT IA mice (E-H) Quantitative qPCR and ELISA analyses showing upregulated inflammatory mediators (TNF-α, NF-κB p65, IL-1β, IL-6, MCP-1, MMP2, and MMP9) in CoW tissues and serum from ARHGEF17^⁻/⁻^ IA mice compared with WT IA. Scale bars = 20 μm. Data are presented as mean ± SEM; n = 6 per group. Statistical significance was determined using one-way ANOVA followed by Tukey’s post hoc test. (*P<0.05, **P<0.01, ***P,****P<0.001).

Notably, these structural defects were substantially more pronounced in ARHGEF17^⁻/⁻^ IA mice than in WT IA mice, indicating that loss of ARHGEF17 markedly aggravates elastic fiber degradation and compromises vascular wall stability during aneurysm formation. (Fig.2A). Elastic and collagen fiber architecture was further examined using Elastica van Gieson (EVG) staining. In WT mice, EVG staining displayed a well-preserved vascular wall structure, with a continuous and compact internal elastic lamina (IEL) adjacent to the endothelium, a uniform tunica media composed of regularly arranged elastic lamellae and smooth muscle cells, and densely organized collagen fibers in the adventitia. In contrast, ARHGEF17^⁻/⁻^ IA mice exhibited extensive vascular wall degradation, characterized by discontinuous, fragmented, or locally absent IEL, disrupted elastic lamellae, and loosely arranged, uneven collagen fibers in the adventitia (Fig. 2B). Importantly, these degenerative changes were considerably more pronounced in ARHGEF17^⁻/⁻^ IA mice than in WT IA counterparts, indicating that loss of ARHGEF17 markedly exacerbates elastic and collagen fiber disruption, leading to severe impairment of vascular wall integrity. ECM remodeling was further evaluated using Picrosirius red (PSR) staining under polarized light. In WT mice, PSR staining revealed dominant orange-red birefringence, reflecting abundant, mature type I collagen fibers arranged in parallel bundles consistent with stable ECM organization. In contrast, ARHGEF17^⁻/⁻^ IA mice exhibited a pronounced shift toward greenish-yellow birefringence, indicating increased deposition of immature type III collagen and disorganized ECM architecture (Fig. 2C). This type I/III collagen imbalance and fragmented fiber arrangement highlight dysfunctional ECM remodeling and compromised mechanical stability of the vascular wall ^[18]^.

ARHGEF17 deficiency also induced VSMC phenotypic switching^[4, 19–21]^. Immunofluorescence staining for α-smooth muscle actin (α-SMA) and osteopontin (OPN) revealed reciprocal expression changes characteristic of a transition from the contractile to the synthetic phenotype. In WT mice, α-SMA showed strong and uniform expression along the tunica media, with circumferentially aligned VSMCs maintaining contractile morphology, whereas OPN expression was minimal. In contrast, ARHGEF17^⁻/⁻^ IA mice exhibited a dramatic reduction in α-SMA intensity and disrupted circumferential organization, accompanied by robust OPN upregulation in both the tunica media and hyperplastic intima (Fig.2D). This α-SMA downregulation/OPN upregulation pattern confirms VSMC phenotypic modulation toward a synthetic, pro-remodeling state associated with increased proliferation and matrix metalloproteinase (MMP) secretion, directly contributing to ECM degradation, tunica media thinning, and vascular wall instability(Fig.2E,F)^[22, 23]^.In parallel with structural damage and VSMC phenotype shifts, inflammatory responses were markedly activated in ARHGEF17^-/-^ IAs mice(Fig.2G,H).In healthy cerebral arteries, inflammation is tightly constrained, but it becomes a dominant driver during IA development^[4, 24, 25]^. ARHGEF17 deficiency triggered the upregulation of multiple inflammatory mediators, as evidenced by quantitative qPCR and ELISA analyses. In the CoW, mRNA levels of TNF-α, NF-κB p65, IL-1β, IL-6, and MCP-1 were significantly upregulated in ARHGEF17^-/-^ IA mice compared to WT mice consistent with transcriptional upregulation, protein levels of these mediators in the CoW were also elevated. Moreover, circulating levels of these pro-inflammatory cytokines in serum were significantly higher in Arhgef17^-/-^ IA mice reflecting systemic inflammatory activation secondary to local vascular injury. This robust inflammatory activation likely contributed to the observed vascular pathologies mediated by NF-κB signaling promotes the release of matrix metalloproteinases ^[26, 27]^, and indeed, MMP2 and MMP9 expression was significantly increased in both the serum and CoW of Arhgef17^-/-^ IAs mice (Fig.2E,F).

Collectively, these findings indicate that ARHGEF17 deficiency orchestrates a pathological cascade involving inflammatory activation, VSMC phenotypic switching, ECM disorganization, and elastic fiber disruption, ultimately driving intracranial aneurysm formation and progression.

### Zebrafish Animal Model Experiments Showed Intracranial Hemorrhage Occurred in The ARHGEF17 Disruption Larvae

To further validate the vascular abnormality in vivo, we constructed ARHGEF 17-deficient zebrafish to mimic the loss of ARHGEF 17 function using morpholino oligonucleotide (MO), in which wild type of AB strain zebrafish embryos was injected with ARHGEF17 Morpholino. Wild-type AB embryos injected with the ARHGEF17-specific morpholino (I9E10-MO) exhibited a striking intracranial hemorrhage phenotype at 72 hours post-fertilization (hpf), as visualized by O-dianisidine staining of erythrocytes (Figure 3A-C). Compared with control morpholino (ctrl-MO) embryos, the I9E10-MO group displayed a significantly higher incidence of intracranial bleeding, indicating that ARHGEF17 disruption compromises cerebrovascular stability. Co-injection of capped ARHGEF17 mRNA markedly rescued the hemorrhagic phenotype, restoring vascular integrity and reducing the hemorrhage rate to near-control levels. In addition, pericardial edema was observed in a subset of I9E10-MO larvae, suggesting systemic circulatory stress.

**Fig. 3.**
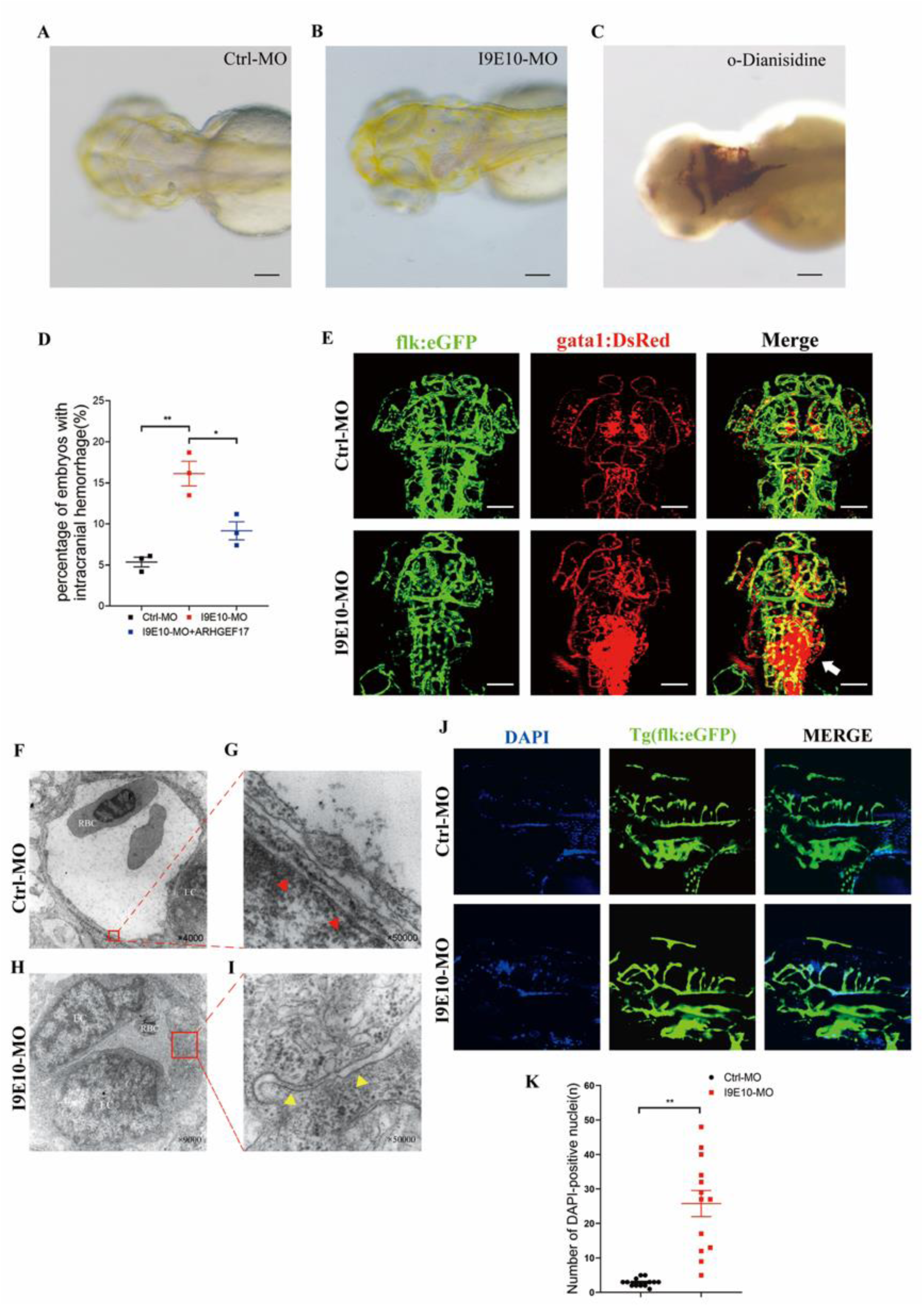
ARHGEF17 Deficiency in Zebrafish Leads to Intracranial Hemorrhage and Impaired Cerebrovascular Integrity (A) Representative bright-field image of wild-type (WT) zebrafish larvae injected with control morpholino (ctrl-MO), showing normal cranial vasculature and absence of hemorrhage at 72 hours hpf. (B) Representative images showing that arhgef17 knockdown with morpholino oligonucleotide (MO) caused intracranial hemorrhage in AB larvae. The images were taken from larvae at 72 hpf. (C) Intracranial hemorrhage were identified with o-Dianisidine staining of erythrocytes. (D) Summary of the intracranial hemorrhage effect of arhgef17 knockdown. The experiments were repeated 3 times and each time > 89 embryos were examined for each group. (E) Double trangenic zebrafish Tg(flk:eGFP; gata1: DsRed) expresse green fluorescent in endothelial cells and red fluorescent in erythrocytes. The images were taken under the confocal microscopy at 72 hpf. Note that gata1+ erythrocytes leaked out from vessels and formed hematomas in the I9E10-MO lavae (white arrow) while erythrocytes remained within vessels in Ctrl-MO lavae. (F-I) Transmission Electron Microscopy (TEM) of intracranial vessels showing endothelial cell swelling, nuclear enlargement, and perivascular edema in I9E10-MO larvae, confirming endothelial activation and structural compromise. (J-K) DAPI leakage assay following injection into the common cardinal vein (CCV) demonstrating impaired vascular barrier integrity. In I9E10-MO larvae, DAPI diffused from intracranial vessels into the surrounding parenchyma within 30 min, indicating increased endothelial permeability (P<0.01 vs ctrl-MO). Data are shown as mean ± SEM and analyzed by one-way ANOVA followed by Tukey’s test. *P<0.05, **P<0.01. Scale bars: 100 μm

(Fig.3D). To further delineate cerebrovascular alterations, we used the Tg(flk:eGFP; gata1: DsRed) double transgenic line, in which endothelial cells and erythrocytes are labeled with eGFP and DsRed, respectively. Confocal imaging revealed extravasation of red blood cells from morphologically intact cerebral vessels, confirming that hemorrhage occurred with gross structural vessel malformation (Fig. 3E).

Transmission electron microscopy (TEM) of cerebral vessels at 72 hpf revealed distinct ultrastructural abnormalities in I9E10-MO larvae, including endothelial cell activation, nuclear swelling, irregular luminal contours, and pronounced perivascular edema (Fig. 3F-I). Tight junctions appeared widened and discontinuous, further supporting the presence of endothelial barrier compromise. To explore the integrity of the intracranial vessels of the larvae without intracranial hemorrhage in I9E10-MO group, we injected DAPI through the CCV of the larvae. After 30 minutes, DAPI could leak into the brain parenchyma beside the blood vessels in I9E10-MO group.

From this, even if the experimental group of larvae without intracranial hemorrhage, the integrity of the intracranial blood vessel wall was also damaged ((Fig.3J, K).

Collectively, these findings demonstrate that ARHGEF17 loss in zebrafish leads to cerebrovascular fragility, endothelial activation, and compromised barrier integrity, phenocopying the intracranial hemorrhage and vascular wall defects observed in ARHGEF17^⁻/⁻^ mice, and supporting a conserved, evolutionarily robust role of ARHGEF17 in maintaining cerebrovascular stability.

### Knockdown of ARHGEF17 Impairs Vascular Endothelial Cell Functions in a Silencing Efficiency-Dependent Manner

To verify the effect of ARHGEF17 on endothelial dysfunction, the lentiviral-mediated ARHGEF17 knockdown in HUVECs yielded efficient and stable gene silencing. qPCR validation confirmed effective silencing of ARHGEF17 mRNA expression in LV-shARHGEF17 group with shRNA-76396 exhibiting higher silencing efficiency (73.31% reduction in mRNA expression) compared to shRNA-76397 and shRNA-76398 (Fig. 4A). A series of cellular function experiments were conducted to investigate the role of ARHGEF17 in ECs. Cell proliferation, assessed via the CCK-8 assay, was significantly attenuated in the LV-shARHGEF17 group compared to the control group. At 24h post-seeding, the absorbance values (OD450) in the LV-shARHGEF17 group were 65.46%,41.54%,42.52% (shRNA-76396, shRNA-76397 and shRNA-76398) lower than those in the control group, with the more efficient shRNA-76396 construct inducing more severe viability reduction. (Fig. 4B). Time-course analysis further demonstrated a progressive exacerbation of proliferation inhibition over 72h, with the viability gap between LV-shARHGEF17 groups and control group widening over time. Consistent with the 24h data, shRNA-76396 exhibited more profound viability impairment than another two LV-shARHGEF17 groups at 72 h, the magnitude of proliferation inhibition was positively correlated with ARHGEF17 silencing efficiency across all time points. (Fig. 4C). Vascular angiogenesis, a key endothelial cell function, was significantly inhibited by ARHGEF17 knockdown in an efficiency-dependent manner. In the tube formation assay, the LV-shARHGEF17 group exhibited severely disrupted vascular-like structure formation compared to the control. Representative images of tube formation assays showed striking reductions in tube-like structure formation and branching in both shRNA-76396 and shRNA-76397 group compared to control group, with the inhibitory effect being more evident in shRNA-76396 (Fig. 4D). Quantitative analysis confirmed significant decreases in both tube junction number and total tube length: shRNA-76396 and shRNA-76397 groups showed 48.67% and 28.00% reductions in tube junctions, respectively, along with 65.70% and 43.37% reductions in total tube length compared to control group (Fig. 4E). These results directly reflect the positive correlation between angiogenesis inhibition and ARHGEF17 silencing efficiency, with the higher-efficiency shRNA-76396 construct inducing more severe impairment of vascular formation capacity. Wound healing assay demonstrated that ARHGEF17 knockdown significantly delayed wound closure. At 24h post-scratch, the wound closure rate in the LV-shARHGEF17 group was lower than that in the control group. Leading-edge cells in the control group displayed well-polarized lamellipodia, whereas ARHGEF17-knockdown cells showed irregular protrusions and reduced cell-to-cell coordination. The weakly silencing group further supporting the efficiency-dependent effect. The effect of ARHGEF17 knockdown on endothelial barrier function was evaluated using trans-endothelial electrical resistance (TEER) measurements, a sensitive indicator of endothelial monolayer integrity. Compared to the control group, the LV-shARHGEF17 group exhibited a significant and progressive reduction in TEER values over the 72-hour monitoring period. Specifically, baseline TEER (measured at 24 hours post-seeding, when monolayers were confluent) was 18% lower in the sh-ARHGEF17 group. By 48 hours, this difference became more pronounced, with TEER values in the knockdown group decreasing to 52% of those observed in controls. As a guanine nucleotide exchange factor that regulates Rho GTPase-mediated actin cytoskeleton remodeling, ARHGEF17 plays a pivotal role in maintaining the structural and functional homeostasis of ECs, a process tightly linked to the organization of filamentous actin (F-actin) and the integrity of vascular endothelial cadherin (VE-Cadherin) to mediated adherens junctions.

**Fig. 4.**
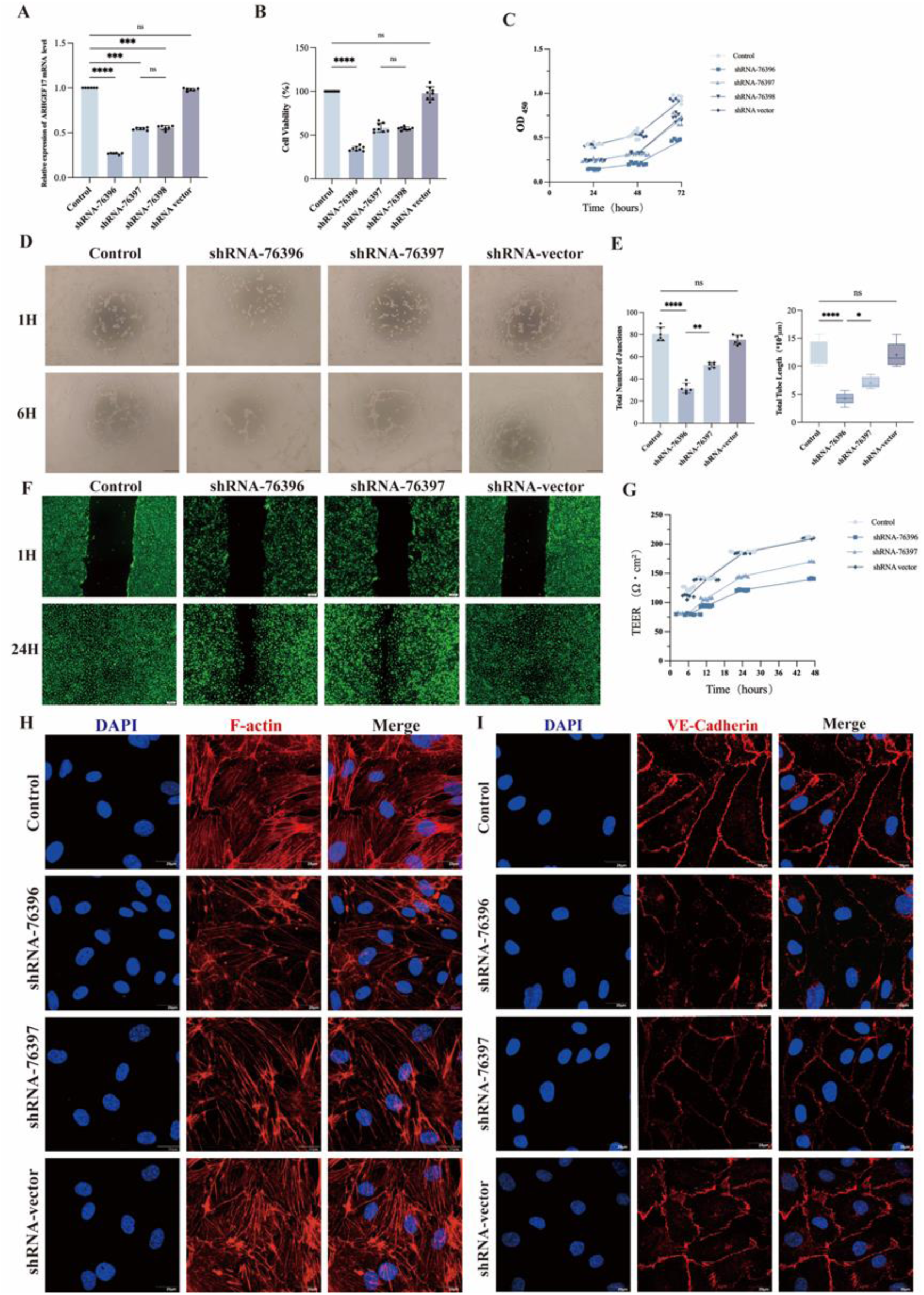
ARHGEF17 Knockdown Impairs Vascular Endothelial Cell Functions in a Silencing Efficiency-Dependent Manner (A) Validation of lentiviral ARHGEF17 knockdown efficiency in HUVECs by quantitative PCR. Among three shRNA constructs (shRNA-76396, shRNA-76397, shRNA-76398), shRNA-76396 achieved the highest silencing efficiency, reducing ARHGEF17 mRNA expression by approximately 73% compared with control. (B) Cell viability measured by CCK-8 assay at 24 h showing a significant decrease in proliferative activity in ARHGEF17-silenced cells, with shRNA-76396 causing the most pronounced inhibition (C) Time-course analysis of cell proliferation over 72 h demonstrating a progressive reduction in viability, consistent with silencing efficiency. (D) Tube formation assay on Matrigel showing that ARHGEF17 knockdown markedly disrupted capillary-like structure formation. Representative images display reduced tube length and branching points in shRNA-76396 and shRNA-76397 groups compared with control. (E) Quantitative analysis of tube junction number and total tube length demonstrating significant reductions in shRNA-76396 and shRNA-76397 groups relative to control. (F) Wound-healing assay showing delayed closure of the scratch area in ARHGEF17 knockdown HUVECs compared with control at 24 h, with impaired lamellipodia formation and reduced directional migration. (G) Trans-Endothelial Electrical Resistance (TEER) measurements showing progressive barrier deterioration in ARHGEF17 knockdown HUVECs. (H-I) Immunofluorescence staining for VE-Cadherin and F-actin demonstrating disruption of adherens junctions and cytoskeletal architecture in ARHGEF17-knockdown HUVECs. VE-Cadherin displayed discontinuous, fragmented borders, while F-actin formed dense stress fibers replacing circumferential actin rings. Scale bars = 20 µm. Data are presented as mean ± SEM from three independent experiments. Statistical significance determined by one-way ANOVA followed by Tukey’s post-hoc test (*P<0.05, **P<0.01, ***P,****P<0.001).

Immunofluorescence staining of ECs further validated the regulatory role of ARHGEF17 in these two key components. In ARHGEF17-knockdown ECs, VE-cadherin displayed discontinuous and fragmented intercellular borders, indicating junctional disassembly, while F-actin reorganized into thick central stress fibers with a concomitant loss of cortical actin rings, reflecting cytoskeletal tension and impaired endothelial stability. These findings indicate that ARHGEF17 knockdown compromises both the basal integrity and dynamic stability of the endothelial barrier, consistent with disrupted intercellular junctional complexes and cytoskeletal organization. Such barrier dysfunction suggests a potential role for ARHGEF17 in maintaining vascular homeostasis through regulation of endothelial permeability.

### ARHGEF17 Deficiency Activates the RhoA/Rock2/MLC Signaling Pathway in Endothelial Cells via Rock2-Dependent Mechanisms

In our prior research, we discovered that ARHGEF17 encodes a Dbl family Rho GEF, which is crucial for activating the Rho family GTPases. ARHGEF17 localizes to the actin cytoskeleton via sequences in the N-terminus and activates RhoA at intercellular junctions. Thus, it is reasonable to assume that ARHGEF17 mutations disrupt RhoA activation, which in turn leads to the destabilization of endothelial junctions in blood vessels, increasing the risk of IAs formation and rupture^[8]^.

To investigate the regulatory role of ARHGEF17 in ECs signaling, we first examined the effects of LV-ARHGEF17 on the RhoA/Rock2/MLC pathway using Western blotting (WB) and GTPase Activation Assay (G-LISA), the latter specifically quantifies the levels of active RhoA (GTP-bound RhoA), total RhoA, and their ratio to accurately reflect RhoA activation status. G-LISA results showed that LV-ARHGEF17 significantly increased the level of active RhoA and the ratio of active-to-total RhoA compared with the control group, with shRNA-76396 exhibiting the most prominent activation effect (Fig.5A-C). Specifically, in shRNA-76396 group, the relative level of active RhoA was 2.93 ± 0.27 (vs. 1.00 ± 0.09 in control), and the active-to-total RhoA ratio was 2.55 ± 0.25 (vs. 1.00 ± 0.05 in control). In contrast, shRNA-76397 and shRNA-76398 induced a milder increase in active RhoA and active-to-total RhoA ratio, while total RhoA expression remained unchanged across all groups. Concomitantly, WB results (Fig.5D) revealed a marked upregulation of Rock2 protein expression in shRNA-76396 group, consistent with the G-LISA data, shRNA-76396 caused the highest Rock2 upregulation. (Fig.5E-F). Given that Rock mediates downstream signaling primarily through phosphorylating its specific substrates, we further detected the phosphorylation levels of myosin light chain 2 (MLC2) and myosin phosphatase target subunit 1 (MYPT1), two classic Rock substrates. The phosphorylation level of MLC at Thr18/Ser19 (p-MLC Thr18/Ser19) was significantly elevated in shRNA-76396 group (1.76 ± 0.22 vs. 1.00 ± 0.08 in control), While the change in total MLC2 expression was not obvious. (Fig.5I, J). We also detected the expression of MLCK,another key regulator of MLC phosphorylation that acts in parallel with the Rho/Rock pathway. WB analysis showed that ARHGEF17 significantly upregulated MLCK protein expression (3.5.92 ± 0.27 vs. 1.00 ± 0.05 in control) (Fig.5K). This result indicates that ARHGEF17 mutation simultaneously activates two core MLC2 regulating kinases, Rock2 and MLCK, which may synergistically contribute to the subsequent increase in MLC2 phosphorylation. For MYPT1, ARHGEF17 mutation specifically increased the phosphorylation of MYPT1 at Thr853(2.15 ± 0.20 vs. 1.00 ± 0.11 in control), and no significant difference in total MYPT1 expression across groups (Fig. 5G.H). These results suggest that ARHGEF17 mutation drives MLC phosphorylation through a dual mechanism, Rock2 activation which direct phosphorylation of MLC and indirect inhibition of myosin phosphatase via p-MYPT1 Thr853 and MLCK upregulation to direct phosphorylation of MLC.

**Fig. 5.**
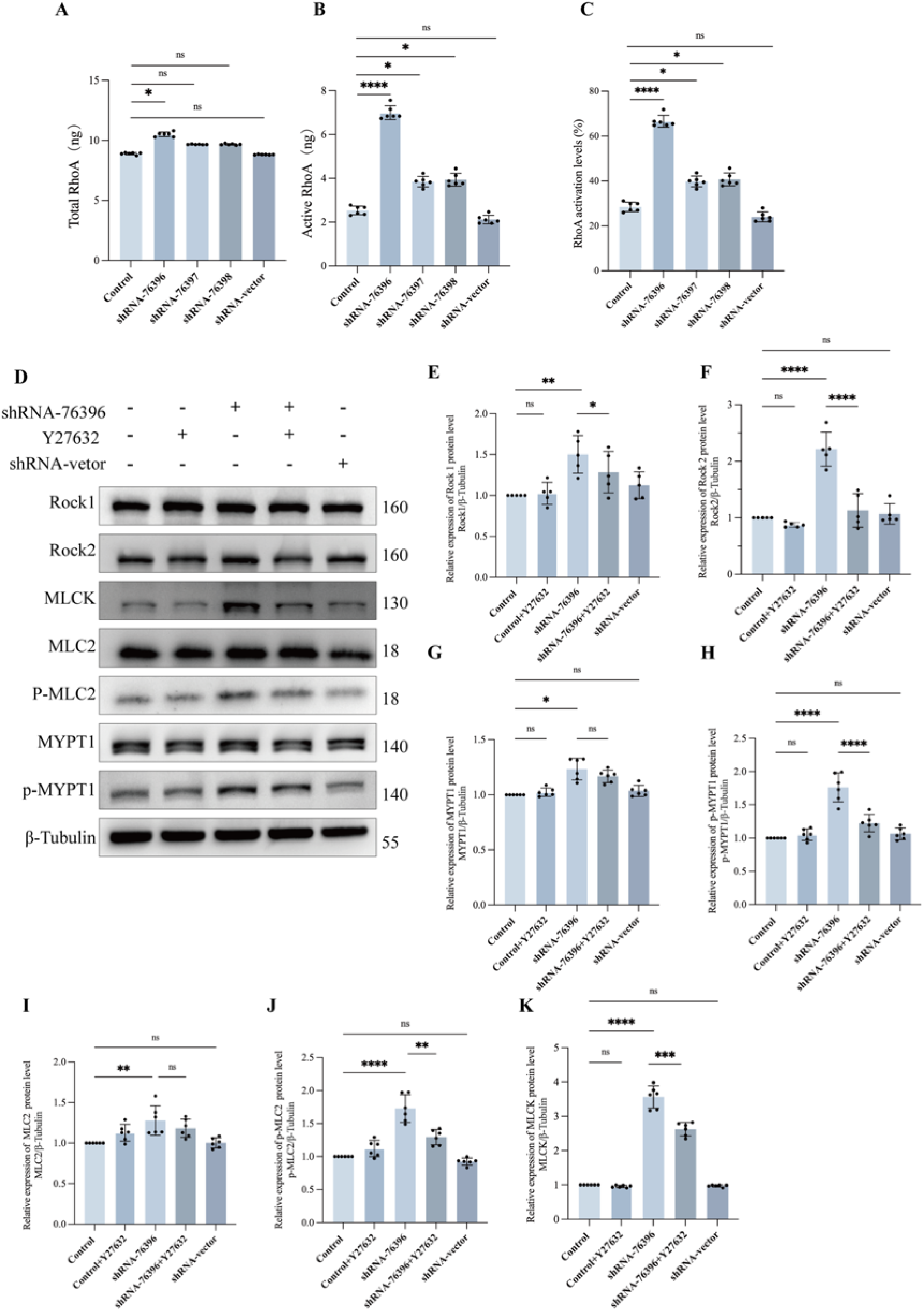
ARHGEF17 Deficiency Activates the RhoA/ROCK2/MLC Signaling Pathway in Endothelial Cells via a ROCK2-Dependent Mechanism (A–C) G-LISA assays measuring RhoA activity in control and ARHGEF17 knockdown HUVECs. Lentiviral knockdown of ARHGEF17 significantly increased the level of active GTP-bound RhoA and the ratio of active-to-total RhoA compared with control cells, with the strongest activation observed in shRNA-76396 cells. (D) Western blot analysis showing protein expression of key components in the RhoA/Rock2/MLC signaling cascade, including Rock1, Rock2, MYPT1, phosphorylated MYPT1 (Thr853), MLC, phosphorylated MLC (Thr18/Ser19), and MLCK. (E-F) Rock1 levels showed a mild increase after ARHGEF17 knockdown, Y-27632 treatment partially suppressed this elevation. ROCK2 expression was significantly increased by approximately 2.4-fold in shRNA-76396 HUVECs compared with controls, and Y-27632 treatment markedly suppressed ROCK2 upregulation, restoring its expression to near-baseline levels. (G-H) Total MYPT1 expression was unaffected by either ARHGEF17 knockdown or Y-27632, confirming that the observed activation occurs via phosphorylation rather than altered protein abundance. p-MYPT1 (Thr853) levels were elevated almost 2.2-fold in shRNA-76396 HUVECs and normalized by Y-27632. (I-K) Total MLC2 levels were unaffected by either treatment. Phosphorylated MLC2 (Thr18/Ser19) was markedly elevated and effectively restored by Y-27632, indicating ROCK2-dependent MLC activation. MLCK expression was significantly upregulated, suggesting cooperative regulation of MLC phosphorylation through both ROCK2 and MLCK pathways. Data are presented as mean ± SEM (n = 6 independent experiments). Statistical significance was determined by one-way ANOVA with Tukey’s post-hoc test (*P < 0.05, **P < 0.01, ***P, ****P < 0.001).

To confirm whether the activation of the RhoA/Rock2/MLC pathway induced by ARHGEF17 is Rock2-dependent, we treated the most effective group shRNA-76396 with the specific Rock inhibitor Y-27632 (10 μM) for 24 h.Y-27632 treatment did not reverse the shRNA-76986 induced increase in active RhoA level or active-to-total RhoA ratio(Supplementary Fig. S3). However, Y-27632 significantly reversed the shRNA-76986-induced upregulation of downstream signaling molecules in this pathway to near-normal levels, including Rock2, phosphorylated MLC at Thr18/Ser19, and phosphorylated MYPT1 at Thr853. Additionally, no significant changes in the expression levels of total RhoA (Supplementary Fig. S3), total MLC and total MYPT1 were observed after Y-27632 treatment. Taken together, these findings demonstrate that ARHGEF17 deficiency activates the RhoA/Rock2/MLC signaling cascade in endothelial cells in a Rock2-dependent manner. Although ARHGEF17 knockdown enhances RhoA activation, inhibition of Rock2 effectively abolishes the downstream phosphorylation of MLC2 and MYPT1, underscoring a pivotal role for Rock2 as the central mediator of RhoA-induced endothelial cytoskeletal remodeling and barrier dysfunction.

### Y-27632 Rescues ARHGEF17 Deficiency-Induced Endothelial Cell Dysfunction by Inhibiting Rock

To further substantiate the hypothesis that the ARHGEF17 mutation compromises ECs functionality through the RhoA/Rock2/MLC signaling cascade, a comprehensive panel of in vitro assays was employed to meticulously assess the restoration of ECs functions, thereby providing conclusive evidence for the role of this signaling pathway in the pathophysiological processes associated with the ARHGEF17 mutation. Tube formation assay was conducted to compare the control group, shRNA-76396 group, and Y-27632 treatment group. After treatment with 10 μM Y-27632 for 24 hours, cells were seeded on Matrigel. Results showed that shRNA-76396 group had a significantly reduced tube formation ability, the Y-27632 treatment rescued this defect, nearly restoring control group like tubulogenic potential (Fig.6A, B). Wound healing assay was employed to determine whether Y-27632 could restore the migratory deficit of ARHGEF17-deficient ECs. At 24 h post-scratch, the wound closure rate of treatment with Y-27632 markedly accelerated the migration of ARHGEF-deficient. This result clearly demonstrates that Y-27632-mediated Rock inhibition effectively ameliorates the impaired migratory capacity of ARHGEF17-deficient ECs (Fig.6C). TEER measurements were performed to quantify the restorative effect of Y-27632 on EC barrier function. Notably, treatment with Y-27632 led to a marked recovery of TEER in ARHGEF17-deficient ECs monolayers, TEER values increased from 125 ±6 Ω·cm² (shRNA-76396) to 176 ±9 Ω·cm² (shRNA-76396+Y-27632) (Fig.6D). To evaluate the impact of Y-27632 on eNOS activation, the phosphorylation levels of eNOS at Ser1177 and Thr495 were analyzed by WB, shRNA-76396 group showed a reduced p-eNOS Ser1177/eNOS ratio and an elevated p-eNOS Thr495/ eNOS ratio compared to control group. In contrast, Y-27632 treatment significantly reversed these abnormalities, the p-eNOS Ser1177/eNOS ratio increased from 0.45 ± 0.15 (shRNA-76396 group) to 0.91 ± 0.16 (shRNA-76396 +Y-27632 group), while the p-eNOS Thr495/ eNOS ratio decreased from 2.95 ± 0.42 (shRNA-76396 group) to 1.88 ± 0.23 (shRNA-76396 +Y-27632 group) (Fig.6E). Immunofluorescence staining was used to assess the restorative effect of Y-27632 on the disruption of endothelial junctions and cytoskeleton induced by ARHGEF17-deficient, with the junctional proteins (ZO-1, VE-Cadherin) and cytoskeletal component F-actin as the key detection targets. shRNA-76396 group showed discontinuous, fragmented distribution of ZO-1 and VE-Cadherin at cell junctions, accompanied by disorganized F-actin, characterized by increased stress fibers and reduced circumferential actin. In contrast, Y-27632 treatment restored the continuous linear distribution of ZO-1 and VE-Cadherin at cell junctions and reorganized F-actin into a regular circumferential belt, indicating that Y-27632 improves junctional and cytoskeletal organization in ARHGEF17-deficient ECs (Fig.6F, G).

**Fig. 6.**
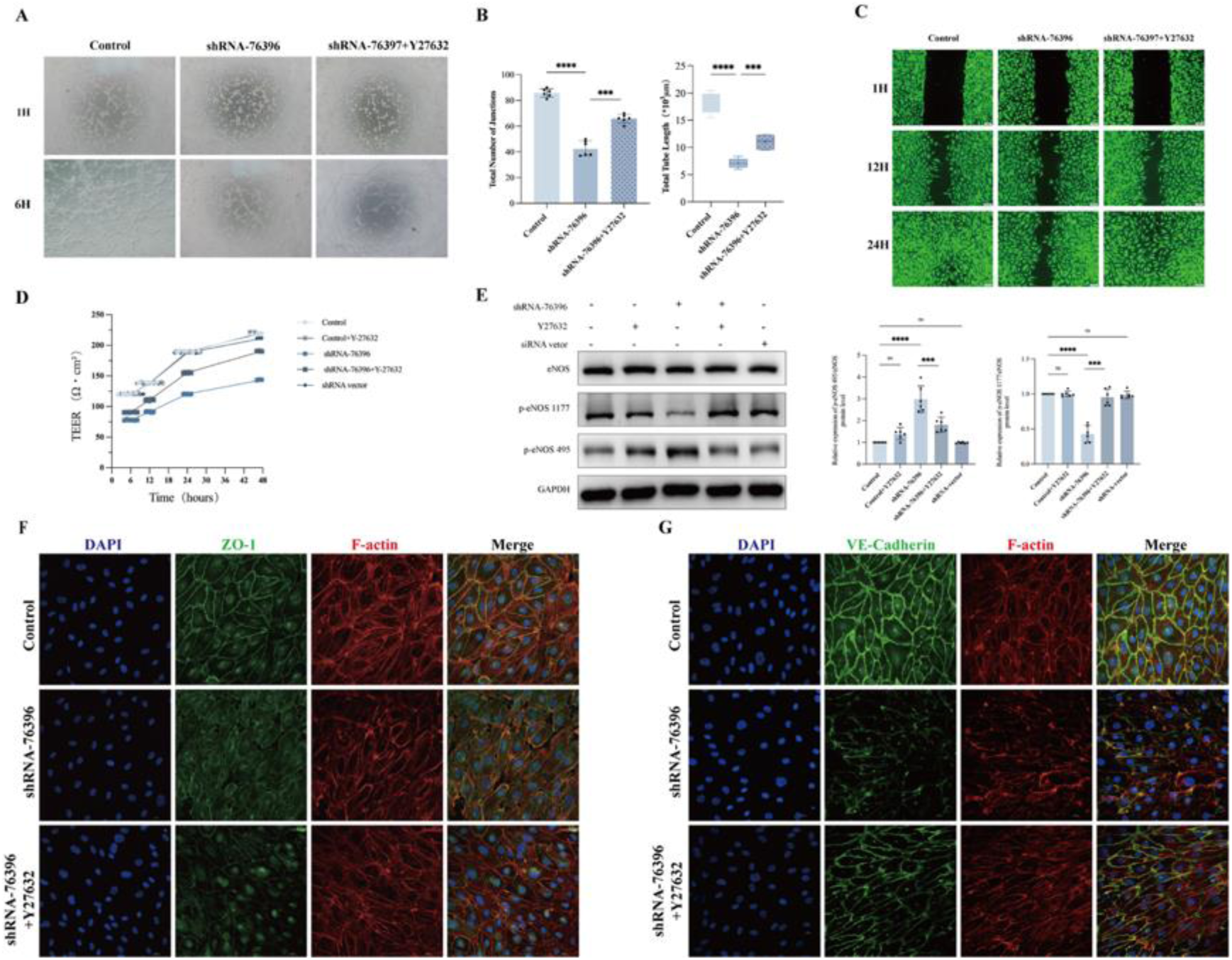
Y-27632 Rescues ARHGEF17 Deficiency–Induced Endothelial Dysfunction via Inhibition of ROCK2 Activity (A–B) Tube formation assay showing impaired angiogenic capacity in ARHGEF17 knockdown HUVECs and its restoration by Y-27632. Quantitative analysis revealed a significant reduction in both tube junction number and total tube length ARHGEF17 knockdown HUVECs, whereas treatment with Y-27632 (10 µM, 24 h) markedly restored tube network complexity. (C) Wound healing assay illustrating the rescue of cell migratory function by Y-27632. The wound closure rate was substantially reduced in shRNA-76396 cells but significantly improved following Y-27632 treatment. Representative images show accelerated closure and polarized lamellipodia formation in the Y-27632 group. (D) Trans endothelial electrical resistance (TEER) measurements indicating restoration of endothelial barrier integrity. ARHGEF17 deficiency decreased TEER values, whereas Y-27632 treatment significantly increased TEER. (E) Western blot analysis of eNOS phosphorylation demonstrating that ARHGEF17 knockdown reduced the activating phosphorylation of eNOS at Ser1177 and increased inhibitory phosphorylation at Thr495. Y-27632 treatment reversed these abnormalities, restoring the p-eNOS Ser1177/eNOS ratio and reducing the p-eNOS Thr495/eNOS ratio. (F–G) Immunofluorescence staining for junctional and cytoskeletal organization (ZO-1, VE-Cadherin, and F-actin). In ARHGEF17-deficient cells, ZO-1 and VE-Cadherin displayed discontinuous, fragmented patterns with increased central stress fibers and loss of cortical actin rings. Y-27632 treatment restored continuous junctional localization of ZO-1 and VE-Cadherin and reorganized F-actin into a circumferential network, indicating recovery of intercellular junction integrity and cytoskeletal stability. Scale bars = 20 µm. Data are presented as mean ± SEM from three independent experiments. Statistical significance determined by one-way ANOVA followed by Tukey’s post-hoc test (*P<0.05, **P<0.01, ***P, ****P<0.001).

Collectively, these findings demonstrate that Rock inhibition by Y-27632 broadly rescues ARHGEF17 deficiency–induced endothelial dysfunction, restoring angiogenesis, migration, barrier integrity, eNOS activation, and cytoskeletal organization. These results provide compelling evidence that ARHGEF17 regulates endothelial homeostasis via the RhoA/Rock2/MLC signaling pathway and highlight Rock inhibition as a potential therapeutic strategy to reverse endothelial impairment associated with ARHGEF17 deficiency.

## Discussion

The present study builds upon our prior whole-genome sequencing efforts that initially identified ARHGEF17 as a candidate gene associated with intracranial aneurysm (IAs) and further validates its functional role in IAs pathogenesis using a comprehensive, multi-model experimental approach^[8]^. By integrating findings from ARHGEF17 knockout (KO) mice, zebrafish models, and lentivirus-mediated ARHGEF17-knockdown cell lines, we provide compelling evidence that dysregulated ARHGEF17 drives IAs formation and rupture by activating the RhoA/Rock2/MLC signaling pathway and subsequent vascular endothelial cells damage, a mechanism that bridges genetic susceptibility to the phenotypic hallmarks of IAs, and fills a critical gap in our understanding of the molecular events underlying this life-threatening vascular disorder.

The RhoA/Rock2/MLC signaling pathway functions as a fundamental regulator of cytoskeletal dynamics, endothelial barrier integrity, and vascular tone, and its aberrant activation constitutes a key pathogenic event following vascular injury^[28, 29]^. Under physiological conditions, transient activation of this axis enables adaptive endothelial responses, such as vasomotor regulation and controlled permeability. However, sustained activation of RhoA/Rock2, induced by mechanical stress, oxidative injury or genetic dysregulation, results in endothelial barrier disruption and vascular inflammation^[30–32]^. RhoA is rapidly converted into its GTP-bound active form, which subsequently activates Rho-associated coiled-coil containing protein kinase 2 (Rock2). Rock2 phosphorylates both myosin light chain (MLC) and myosin phosphatase target subunit 1 (MYPT1), leading to actomyosin contractility and stress fiber formation. This hypercontractile state disrupts endothelial junctions anchored by VE-cadherin, increases paracellular permeability, and compromises endothelial barrier integrity, ultimately promoting vascular leakage and inflammatory cell infiltration^[33–38]^. This dual mechanism maintains MLC in a hyperphosphorylated state, promoting actomyosin contractility, stress fiber formation, and increased cellular tension, forms a crucial molecular link between RhoA/Rock2 activation and vascular injury^[39]^.

In our study, ARHGEF17 deficiency led to enhanced Rock2 expression and markedly increased phosphorylation of both MYPT1 and MLC, consistent with excessive actomyosin contractility. Inhibition of Rock2 with Y-27632 normalized MYPT1 and MLC phosphorylation and restored endothelial barrier function, confirming that Rock2 acts as a key molecular switch mediating the deleterious effects of RhoA activation. Similar findings have been reported in models of atherosclerosis, cerebral vasospasm, and hypertensive vascular remodeling, where RhoA/ROCK2-driven MLC hyperphosphorylation induces vascular stiffening, smooth muscle dedifferentiation, and impaired vasodilation^[39–42]^.

A key strength of this study lies in the cross-species and multi-scale validation of ARHGEF17’s function. Utilizing a well-established ARHGEF17^⁻/⁻^ IAs model mice, we systematically demonstrated a statistically significant elevation in both IA incidence and rupture rate, concomitant with overt vascular wall deterioration that was spatially restricted to the CoW—the anatomical locus that mirrors the primary site of IA development. Histopathological and molecular analyses further delineated that this vascular damage was characterized by hallmark features of pathological vascular remodeling, manifested in two distinct yet interconnected aspects. Severe vascular endothelial cells structural damage and marked upregulation in the secretion and enzymatic activity of matrix metalloproteinases, specifically MMP-2 and MMP-9. The increased MMP levels in ARHGEF17^-/-^ IAs model mice directly contributed to breakdown of the vascular wall’s structural integrity, as evidenced by reduced collagen content and disorganized elastic fibers in the tunica media of CoW. These findings establish a direct link between ARHGEF17 mutation and IA-associated vascular pathology, particularly at the anatomically relevant CoW. Complementary experiments in zebrafish, an ideal model for studying vascular development and genetic perturbations, further confirmed that introduction of ARHGEF17 mutation led to ECs disorganization, impaired vascular barrier function, and abnormal cerebral vascular remodeling, recapitulating the pathogenic phenotype seen in mice and ruling out species-specific artifacts. At the cellular level, lentivirus-mediated ARHGEF17 knockdown in human umbilical vein endothelial cells triggered robust activation of the RhoA/Rock2/MLC signal pathway, which in turn induced prominent ECs dysfunction. Significantly impaired directional cell migration, markedly reduced tube formation capacity, obvious cytoskeletal disruption, compromised barrier function, and notably enhanced secretion of matrix metalloproteinases, a family of enzymes that mediate extracellular matrix degradation and further contribute to endothelial dysfunction and tissue remodeling disorders. Collectively, these data establish a causal chain ARHGEF 17 mutation, RhoA/Rock2 pathway activation damage, IA formation/rupture and validate ARHGEF17 as a functional driver of IA, rather than a passive genetic marker.

Our findings align with and extend the growing body of literature implicating Rho GTPase signaling in vascular homeostasis and IAs pathogenesis. RhoA, a central member of the Rho GTPase family, regulates cytoskeletal dynamics, cell adhesion, and endothelial barrier function—processes that are dysregulated in IA lesions.

Rock2, a downstream effector of RhoA, has been shown to promote vascular smooth muscle cell dedifferentiation and extracellular matrix degradation in AAA or TAA[43, 44]. However, its role in IA, particularly in ECs, remains less defined. Here, we demonstrate that Rock2 activation is critical for ARHGEF17 mediated ECs damage, pharmacological inhibition of Rock2 Y-27632 in ARHGEF17 knockdown HUVECs reversed ECs dysfunction, restoring cell migration and tube formation while reducing inflammation. This observation not only identifies Rock2 as a key mediator of ARHGEF17’s pathogenic effects but also highlights the RhoA/Rock2/MLC axis as a potential therapeutic target for IA, an insight that complements recent preclinical studies showing Rock inhibitors reduce aneurysm growth.

Notably, our study also addresses a longstanding question in IA genetics, how genetic variants associated with IAs translate into functional changes that drive disease. While genome-wide association studies (GWAS) have identified numerous IA susceptibility loci, few have been functionally validated, limiting our ability to develop mechanism-based therapies^[5–7, 29, 45]^. By showing that ARHGEF17 mutation disrupts RhoA/Rock2/MLC signaling pathway across multiple models, we provide a framework for interpreting ARHGEF17 genetic variants. For example, variants that increase ARHGEF17 expression or enhance its GEF activity may predispose individuals to IA by amplifying RhoA/Rock2 mediated ECs damage, ARHGEF17 may interact with other Rho GTPases (Rac1, Cdc42) or signaling cascades (MAPK, PI3K/Akt) that contribute to IA pathogenesis, investigating these alternative pathways will provide a more comprehensive view of ARHGEF17’s role in IA ^[46, 47]^.Future studies focusing on the functional impact of specific ARHGEF17 variants, single-nucleotide polymorphisms, copy number variations in patient-derived endothelial cells will further refine this model and may enable personalized risk stratification for IA.

Despite these strengths, our study has several limitations that warrant consideration. First, while our animal models (mice, zebrafish) recapitulate key features of IA, they do not fully mirror the complexity of human IA, which is influenced by aging, hypertension, smoking, and other comorbidities. For instance, the mouse model of IA typically involves induced hypertension or vascular injury, which may not capture the “spontaneous” IA development seen in humans with genetic predisposition. Second, our cell culture experiments used HUVECs, a widely used but non-brain-specific endothelial cell type, future studies using HBAECs or brain organoid models will help confirm whether our findings are specific to cerebral vasculature. A complete understanding of ARHGEF17 mutation dynamics in IAs will necessitate future studies utilizing endothelial lineage tracing techniques or longitudinal single-cell RNA sequencing of intracranial aneurysm tissues. These approaches would allow comprehensive monitoring of endothelial phenotypic transitions during disease evolution. Despite these limitations, our current findings suggest that therapeutic strategies targeting RhoA/Rock2/MLC may offer promising opportunities for early IA intervention.

In conclusion, the present study validates ARHGEF17 as a functional driver of intracranial aneurysm formation and rupture, mediated by activation of the RhoA/Rock2/MLC signaling pathway and subsequent vascular endothelial cell damage. Our multi-model approach confirms the conserved role of this mechanism across species and scales, bridges genetic susceptibility to pathogenic phenotypes, and identifies the RhoA/Rock2/MLC axis as a promising therapeutic target. These findings not only advance our understanding of IA biology but also lay the groundwork for future studies aimed at developing targeted therapies to prevent IA formation and rupture, an unmet clinical need given the high morbidity and mortality associated with this disease.

## Acknowledgments

This work was supported by and National Natural Science Foundation of China (82271302,81571144,82201530). The authors would like to kindly thank Neuroendocrinology Laboratory, Tianjin Institute of Neurology, Tianjin Medical University General Hospital.

## Author contributions

Jian Li * and Hao Zhang * contributed equally.

Conception and design of the research: Jian Li, Hao Zhang; acquisition of data: Jian Li, Hao Zhang, Bangyue Wang; analysis and interpretation of the data: Jian Li, Chao Peng; statistical analysis: Jian Li, Chao Peng; supervising the experiments: Xinyu Yang; drafting the manuscript: Jian Li,Hao Zhang.

## Supplemental Material

### Supplemental Methods Methods

#### Mice

C57BL/6J mice (male, 8-10 weeks old) were sourced from Beijing Vital River Laboratory Animal Technology Co., Ltd. ARHGEF17 knockout (ARHGEF17^⁻/⁻^) mice were generated via a CRISPR/Cas9-mediated gene editing strategy designed in our laboratory and executed by the Shanghai Model Organisms Center, Inc. (Shanghai, China). Homozygous mutants were maintained on a C57BL/6J background and used for experiments after at least six generations of backcrossing. Mice were group-housed in ventilated cages within a specific pathogen-free (SPF) facility and maintained on a 12-hour light/dark cycle. They were provided autoclaved standard laboratory chow and water ad libitum. All animal procedures were conducted in accordance with the Guide for the Care and Use of Laboratory Animals published by the US National Institutes of Health (NIH Publication, 8th edition, 2011) and were approved by the Animal Care and Use Committee at the Institute of Laboratory Animal Science, Tianjin medical university general hospital.

#### Genotype Identification

Genotyping of ARHGEF17 knockout (ARHGEF17^⁻/⁻^) mice was performed using genomic DNA extracted from tail or ear punch tissue collected at weaning (2–3 weeks of age). Cut mouse tissue and place it in a 200 µl centrifuge tube with a numbered label, then close the lid. For each mouse, prepare the reaction solution according to the following: 0.5 µl KAPA Express Extract Enzyme, 2.5 µl 10X KAPA Express Extract Buffer, and 22 µl DEPC Water. Add the solution to the 200 µl centrifuge tube containing the mouse tissue. Place the centrifuge tubes in the reverse transcription instrument and extract DNA following the protocol in Table.S4, PCR products were separated by electrophoresis on 1%(P1P2) and 3%(P3P4) agarose gels containing SYBR™ Safe DNA Gel Stain and visualized under UV illumination. Wild type. P1 and P2 yield a single 5.4 kb fragment, P3 and P4 yield a 500 bp fragment.Heterozygote.P1 and P2 yield 5.4 kb and 1.5 kb fragments (the 5.4 kb fragment may not be obtained due to primer competition in PCR), P3 and P4 yield a 500 bp fragment.Homozygote.P1 and P2 yield a single 1.5 kb fragment, P3 and P4 do not yield a 500 bp fragment.

#### Zebrafish Strain Maintenance

The zebrafish strain Tg(flk: eGFP; gate1: DsRed), AB, was used. All experiments involving zebrafish were approved by the Tianjin Medical University General Hospital Animal Ethics Committee.

According to previous research, zebrafish were raised under standard conditions with a 10-h dark, 14-h light cycle and at 28.5°C. All work involving zebrafish and mice was reviewed and approved by Animal Ethics Committee of Tianjin Medical University General Hospital.

#### Intracranial Aneurysm (IA) Model

Intracranial aneurysms were induced in mice using a well-established model combining pharmacologically induced hypertension with stereotactic elastase injection, as described previously with minor modifications. Briefly, mice (8–10 weeks old, 22–28 g) were anesthetized with 2% isoflurane in 70% N₂O/30% O₂, and core temperature was maintained at 37 ± 0.5 °C throughout the procedure. To induce hypertension, a 50 mg deoxycorticosterone acetate (DOCA) pellet was implanted subcutaneously in the dorsal region under aseptic conditions. vascular injury was induced by stereotactic injection of 17.5 mU elastase dissolved in 2.5 µL sterile PBS into the right basal cistern.

The stereotactic coordinates relative to the bregma were posterior 2.5 mm, lateral 1.0 mm, and depth 5.0 mm. Elastase was infused at a rate of 0.25 µL/min using a microinjection pump over 10 min, and the needle was left in place for an additional 10 min to prevent reflux. Control animals received equal volumes of sterile PBS. Mice were given 1% NaCl in drinking water ad libitum to maintain mineralocorticoid-induced hypertension.

#### Histological Examination

At the experimental endpoint, mice were deeply anesthetized and perfused transcardially with phosphate-buffered saline (PBS, pH 7.4) followed by 4% paraformaldehyde (PFA). The brains were removed, post-fixed in 4% PFA overnight at 4 °C, and the Circle of Willis (CoW) region was carefully dissected under a stereomicroscope Tissues were dehydrated through a graded ethanol series, embedded in paraffin, and sectioned at 8 µm thickness using a rotary microtome. For structural assessment, sections were subjected to multiple histological staining. Victoria Blue staining was performed to visualize elastic fibers. Elastic fibers appeared as blue linear structures, allowing the assessment of fiber continuity, orientation, and integrity. Elastica van Gieson (EVG) staining was conducted to simultaneously distinguish elastic fibers (black or dark blue) and collagen fibers (red), enabling evaluation of internal elastic lamina (IEL) continuity and medial organization. Picrosirius Red (PSR) staining was used to assess collagen composition and remodeling. Under polarized light, type I collagen exhibited red–orange birefringence, whereas type III collagen showed greenish-yellow birefringence, permitting the quantification of collagen subtype ratios and extracellular matrix (ECM) remodeling. For immunofluorescence staining, sections were deparaffinized, rehydrated, and subjected to antigen retrieval in 10 mM citrate buffer (pH 6.0) at 95 °C for 20 min. After blocking with 5% normal goat serum for 1 h, sections were incubated overnight at 4 °C with primary antibodies against α-smooth muscle actin, osteopontin, and CD31. Alexa Fluor conjugated secondary antibodies were applied for 1 h at room temperature, followed by nuclear counterstaining with DAPI. Images were captured using a fluorescence microscope or confocal system and analyzed using ImageJ to quantify mean fluorescence intensity and positive area fraction.

#### Cell culture and treatment of Human Umbilical Vein Endothelial Cells (HUVECs)

Primary Human Umbilical Vein Endothelial Cells (HUVECs) were acquired from CTCC and cultured in endothelial cell medium (ECM, ScienCell) supplemented with 2% Endothelial Cell Growth Supplement (ECGS, ScienCell) and 10% fetal bovine serum (FBS, ScienCell). Cells were maintained in a humidified atmosphere at 37℃ with 5% CO2.Constructed ARHGEF 17 Lentivirus was transfected according to the manufacturer’s protocol. The complete medium was replaced after 24 h of transfection. Rock inhibitor Y-27632 (Dihydrochloride) was resuspend in water. stock solution was diluted into culture medium immediately before use. The medium containing the agents was replaced daily to ensure continuous drug action. Post Lentivirus infection, HUVECs were starved for 24 h and stimulated with 10umol of Y-27632 (Dihydrochloride) for 24h.

#### Western blot

Whole-cell lysates were lysed in ice-cold RIPA lysis buffer (HY-K1001, MCE) supplemented with Protease Inhibitor Cocktail (HY-K0010),Phosphatase Inhibitor Cocktails (HY-K0021、HY-K0022、 HY-K0023) for 20 minutes. Samples were clarified by centrifugation at 13,000 rpm for 20 minutes and protein content was measured using the BCA protein assay kit (Thermo Fisher Scientific). Equal amounts of total proteins (20 μg) were resolved on 6-15% SDS-PAGE gels and transferred onto PVDF membranes (0.45 μm or 0.22μm, Millipore).Membranes were blocked in 5% non-fat dry milk or 5% BSA solution for 2 hours and then incubated overnight at 4 °C with the primary antibody solution, Details of all primary antibodies used were provided in the Table S1.The membranes were then washed with TBS-T,Subsequent to the appropriate horseradish peroxidase-conjugated secondary antibody for 1 hour at room temperature. Signals were detected using the Immobilon Western Chemiluminescent HRP Substrate. Uncropped and unprocessed scans of all the blots in the paper are available in the <u>Supplementary Data</u>.

#### Tube Formation Assay

Matrigel was thawed overnight at 4 °C, kept on ice, and dispensed at 50 µL per well into prechilled 96-well plates. The plates were incubated at 37 °C for 30–45 min to allow polymerization.

HUVECs were serum-starved before the assay. Cells were seeded onto Matrigel-coated wells at a density of 1.5–2.0 × 10⁴ cells/well in 100 µL medium. Where indicated, cells were pretreated for 24hours with the ROCK inhibitor Y-27632 (10 µM) or vehicle control prior to seeding. The plates were incubated at 37 °C with 5% CO₂, and tube-like structures were visualized at 1, 4, and 6 h using a phase-contrast inverted microscope (Olympus IX73). Three random fields per well were imaged under identical conditions. Quantitative analysis was performed using ImageJ (NIH) with the Angiogenesis Analyzer plugin by a blinded investigator.

#### Wound healing analysis

HUVECs with different treatments were suspended in complete medium at a concentration of 7 × 10^5^ cells/ml, and 70 μl of cell suspension were applied into each well of the ibidi Culture-Insert 2 Well system (Ibidi, No.80209 Germany) positioned in a 6-well. When the cell density reached more than 90% confluence, the culture insert was gently removed using sterile tweezers. Meanwhile, HUVEC was replaced with serum free medium. HUVECs were stained with Calcein-AM, and Images were acquired using an inverted microscope at 0 h, 12 h and 24 h after scratch. The percentage of the reduced area was measured using NIH ImageJ software (version 1.52a) and reported as a percentage. Numbers of migrated cells were assessed under light microscopy by counting six fields.

#### Trans endothelial electric resistance (TEER) detection

TEER measures the electrical resistance across the endothelial monolayer, reflecting the integrity and permeability of the blood vessel, with lower values indicating barrier dysfunction. HUVECs were seeded in a 12-well Transwell culture plate. After completing the drug treatment, the resistance of each well was measured using an electrical resistance meter. During the measurement, the longer end of the electrode from the resistance meter was in contact with the bottom of the culture plate, while the shorter end was positioned near the bottom of the chamber. Three different locations within each chamber were selected for measurement, and the average value was taken as the actual TEER measured value. The resistance values were expressed in Ω·cm². The actual TEER was calculated by subtracting the blank value from the measured value.

#### ELISA

Mice were anesthetized with isoflurane, and blood was obtained by cardiac puncture into additive-free tubes. After clotting at room temperature for 30 mins, samples were centrifuged (2,000 × g, 10 min, 4 °C), the supernatant was recentrifuged (10,000 × g, 5 min) to remove residual cells. The CoW was micro dissected on ice, briefly rinsed in ice-cold PBS to remove blood, blotted, snap-frozen in liquid nitrogen. Lysates were incubated on ice for 30 min with gentle agitation to allow efficient protein extraction, then centrifuged at 12,000 × g for 10 min at 4 °C to remove debris. The resulting supernatants were collected and transferred to pre-chilled tubes. Levels of MMP-2, MMP-9, TNF-α, IL-6, IL-1β were measured in serum; MMP-2, MMP-9, NF-κB p65 were assayed in CoW lysates, using mouse-specific sandwich ELISA kits. Because NF-κB p65 is a nuclear transcription factor, it was quantified only in tissue lysates, not in serum. All steps followed the manufacturer’s instructions.

#### Measurement of active RhoA-GTP levels

RhoA activity in HUVECs was detected by G-LISA (BK124, Cytoskeleton, US) according to the manufacturer’s instructions, and normalized to total RhoA levels, which were measured using the Total RhoA ELISA kit (BK150, Cytoskeleton) according to the manufacturer’s instructions.

---

**Table S1.** Antibodies for WB and IF.

| Target antigen | Vendor or Source | Catalog # | Working concentration | Lot # (preferred but not required) | Persistent ID / URL |
| --- | --- | --- | --- | --- | --- |
| eNOS Antibody | CST | #9572 | 1:1000(WB) |  |  |
| Phospho-eNOS (Ser1177) Antibody | CST | #9571 | 1:1000(WB) |  |  |
| Phospho-eNOS (Thr495) Antibody | CST | #9574 | 1:1000(WB) |  |  |
| Anti-MMP9 antibody [RM1020] | abcam | ab283575 | 1:1000(WB) |  |  |
| Anti-MMP2 antibody [EPR1184] | abcam | ab92536 | 1:1000(WB) |  |  |
| Anti-ROCK1 antibody [EPR638Y] | abcam | Ab134181 | 1:1000(WB) |  |  |
| ROCK2 (E5T5P) Rabbit | CST | #47012 | 1:1000(WB) |  |  |
| MYPT1 Antibody | CST | #2634 | 1:1000(WB) |  |  |
| Phospho-MYPT1 (Thr853) Antibody | CST | #4563 | 1:1000(WB) |  |  |
| Myosin Light Chain 2/MLC-2V Polyclonal antibody | Proteintech | 10906-1-AP | 1:1000(WB) |  |  |
| Phospho-Myosin Light Chain 2 (Thr18/Ser19) Polyclonal antibody | Proteintech | 29504-1-AP | 1:1000(WB) |  |  |
| MYLK Polyclonal antibody | Proteintech | 21642-1-AP | 1:1000(WB) |  |  |
| β-Actin (13E5) Rabbit mAb | CST | #4970 | 1:1000(WB) | | |
| β-Tubulin Antibody | CST | #2146 | 1:1000(WB) | | |
| GAPDH (14C10) Rabbit mAb | CST | #2118 | 1:1000(WB) |  |  |
| Peroxidase-conjugated AffiniPure® Goat Anti-Mouse IgG (H+L) | Jackson | 115-035-003 | 1:5000(WB) |  |  |
| Peroxidase-conjugated AffiniPure® Goat Anti-Rabbit IgG (H+L) | Jackson | 111-035-003 | 1:5000(WB) |  |  |
| Donkey anti-Goat IgG (H+L) Secondary Antibody, HRP | Invitrogen | A15999 | 1:5000(WB) |  |  |
| Anti-alpha smooth muscle Actin antibody [EPR5368] | abcam | Ab124964 | 1:200(IF) |  |  |
| osteopontin/OPN/SPP1 antibody (AKm2A1): | Santa Cruz | sc-21742 | 1:100(IF) |  |  |
| CD31/PECAM-1 Antibody | R&D Systems | AF3628 | 1:100(IF) |  |  |
| ZO-1 Monoclonal Antibody (ZO1-1A12) | Invitrogen | 33-9100 | 1:200(IF) |  |  |
| VE-cadherin Antibody (F-8) | Santa Cruz | sc-9989 | 1:100(IF) |  |  |
| Donkey anti-Mouse IgG (H+L) Highly Cross-Adsorbed Secondary Antibody, Alexa Fluor™ 488 | Invitrogen | A-21202 | 1:400(IF) |  |  |
| Donkey anti-Rabbit IgG (H+L) Highly Cross-Adsorbed Secondary Antibody, Alexa Fluor™ 488 | Invitrogen | A-21206 | 1:400(IF) |  |  |
| Donkey anti-Goat IgG (H+L) Cross-Adsorbed Secondary Antibody, Alexa Fluor™ 488 | Invitrogen | A-11055 | 1:400(IF) |  |  |
| Donkey anti-Rabbit IgG (H+L) Highly Cross-Adsorbed Secondary Antibody, Alexa Fluor™ 594 | Invitrogen | A-21207 | 1:400(IF) |  |  |
| Phalloidin-iFluor 647 Reagent | abcam | ab176759 | 1:1000(IF) |  |  |

**Table S2.**
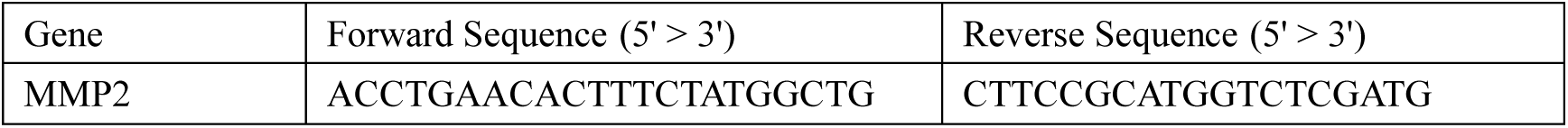
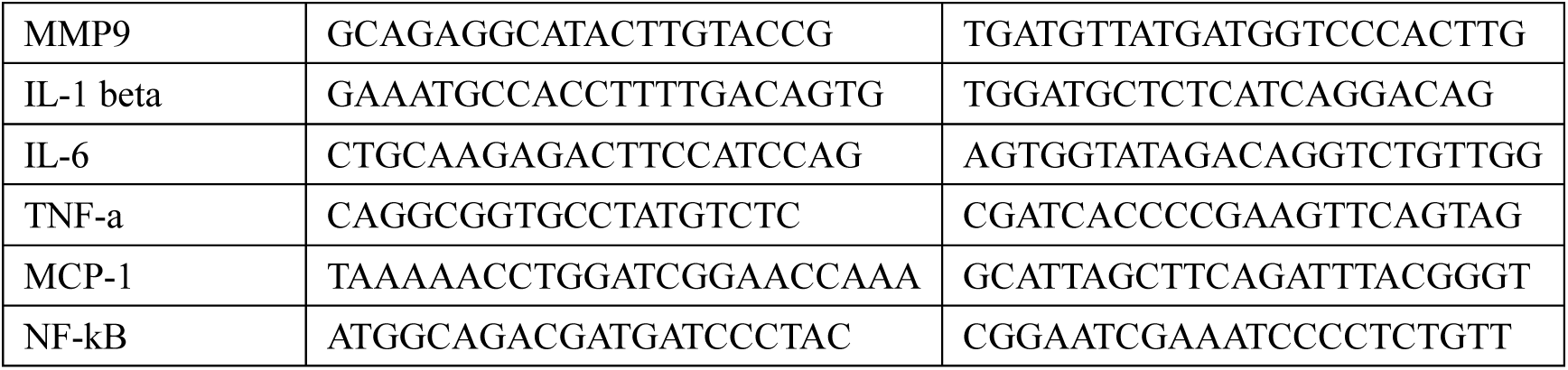
List of primer sequences used for qPCR.

**Table S3.**
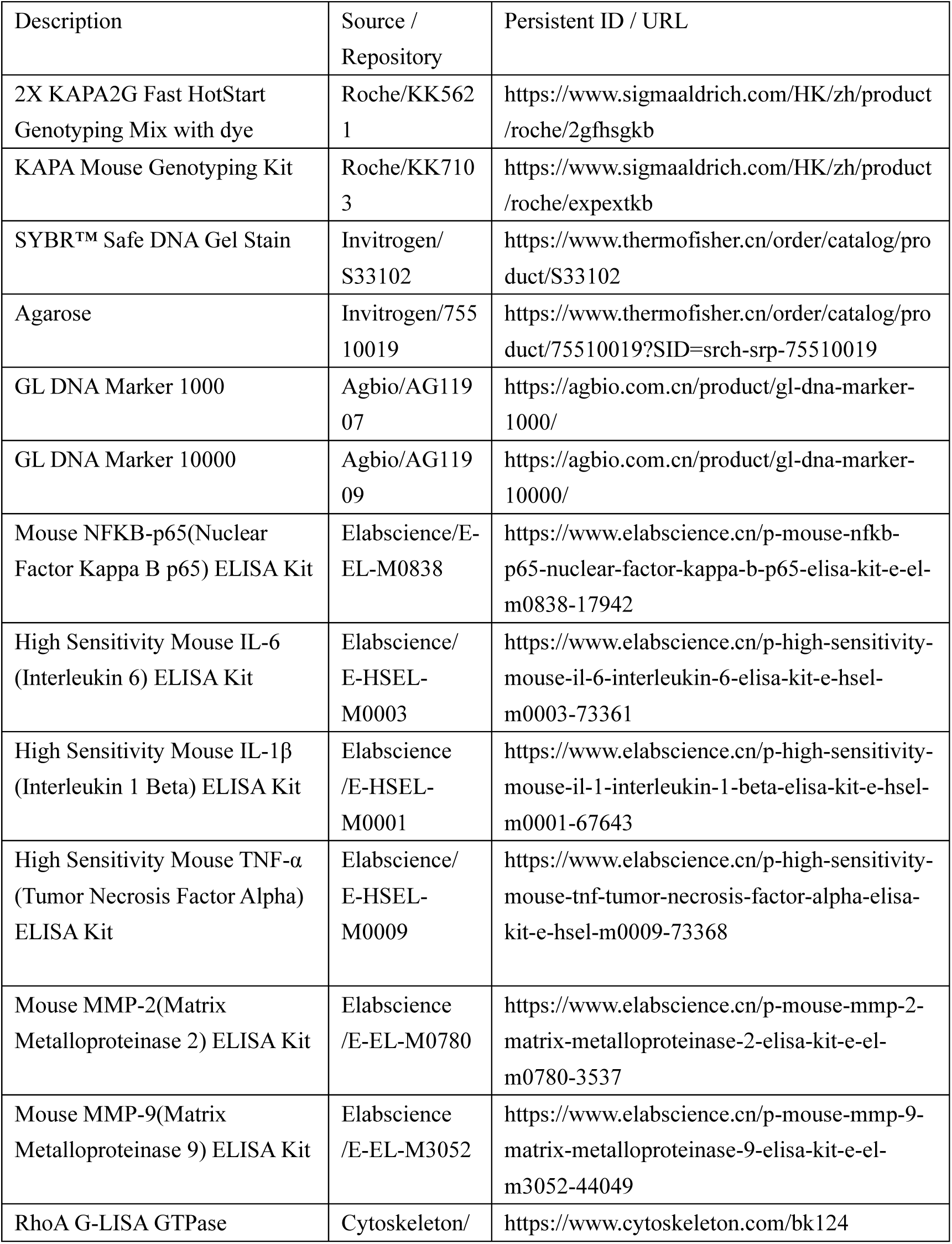
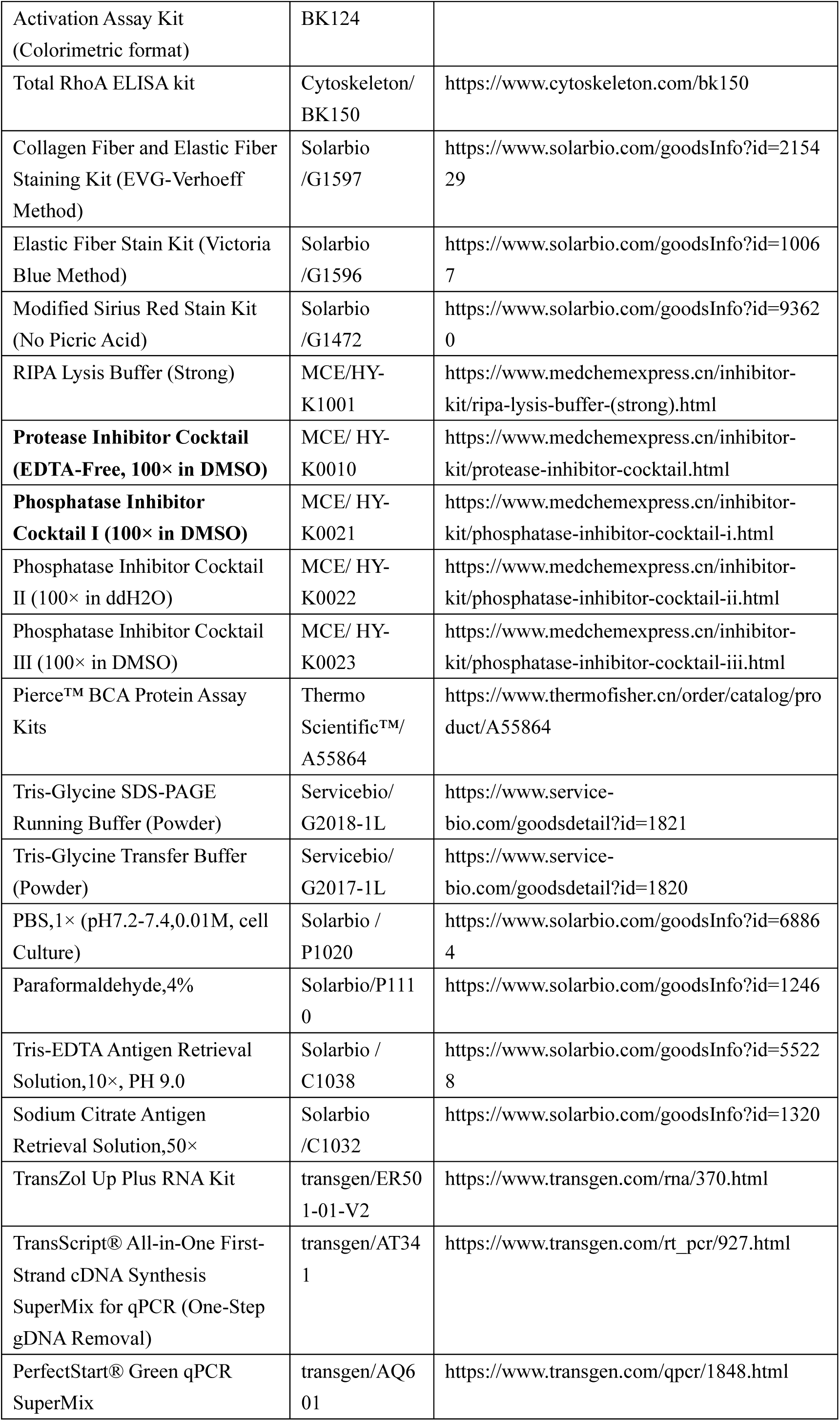
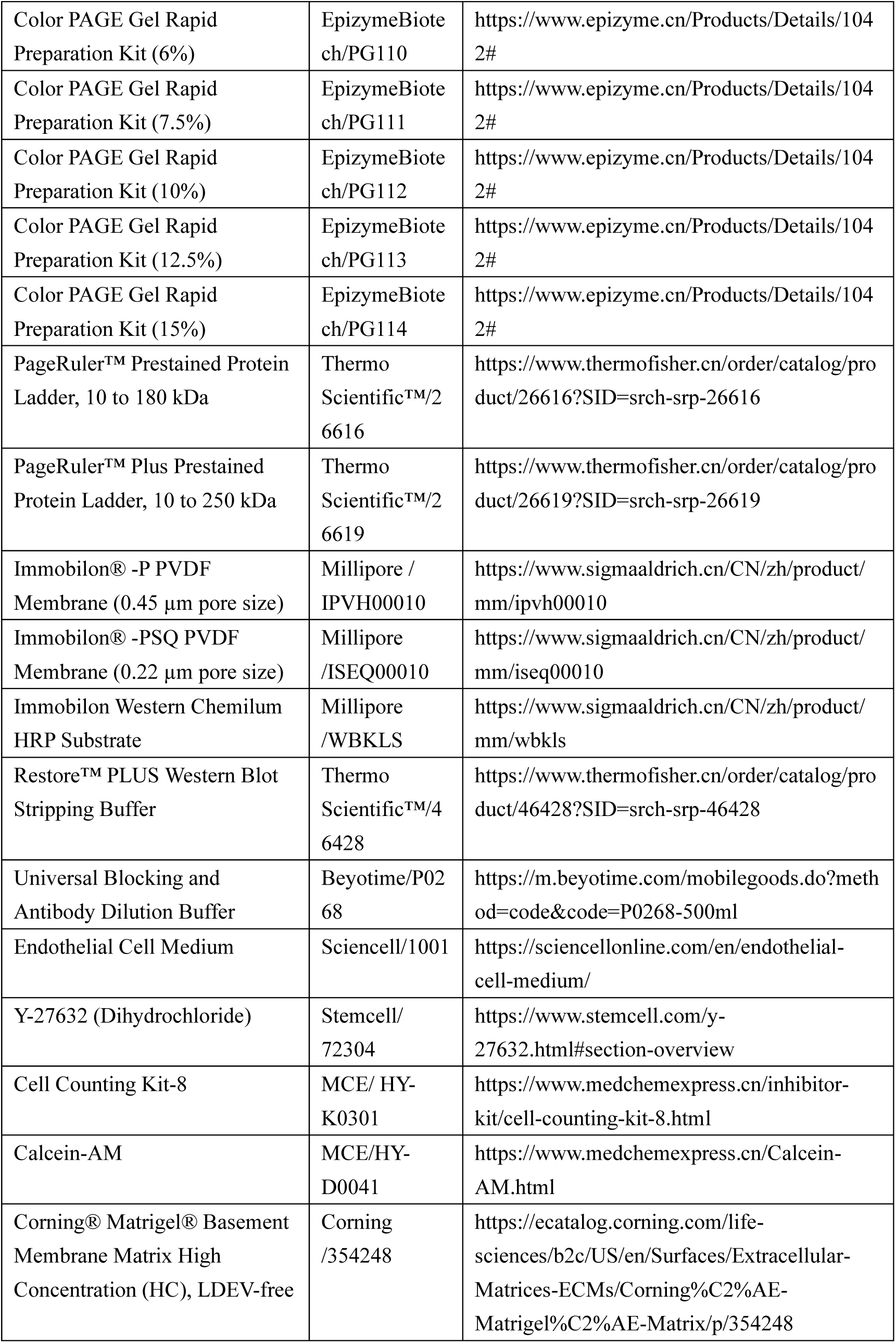
List of Chemical Reagents.

**Table S4.**
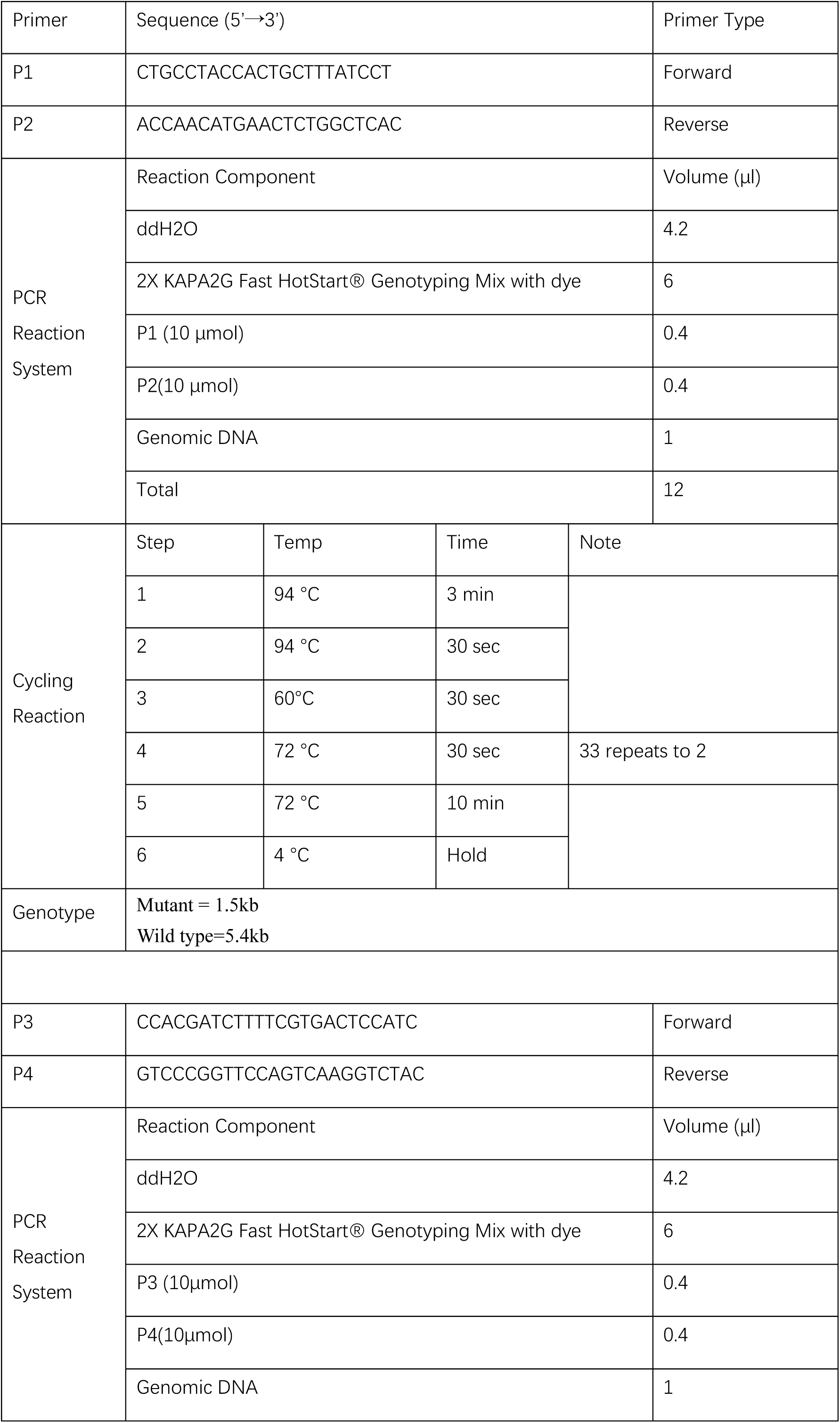
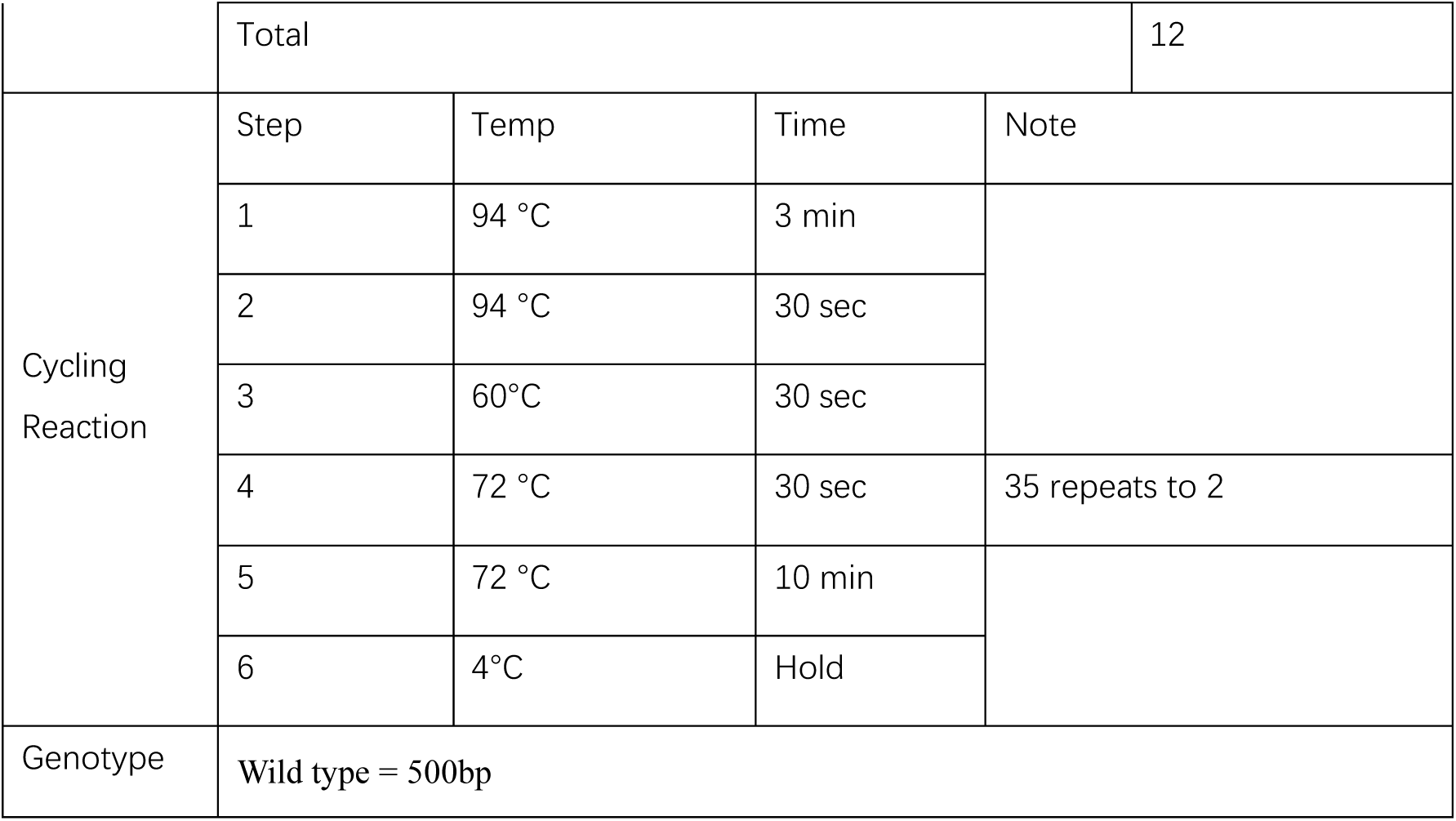
Step of PCR for genotype identification.

**Fig. S1.**
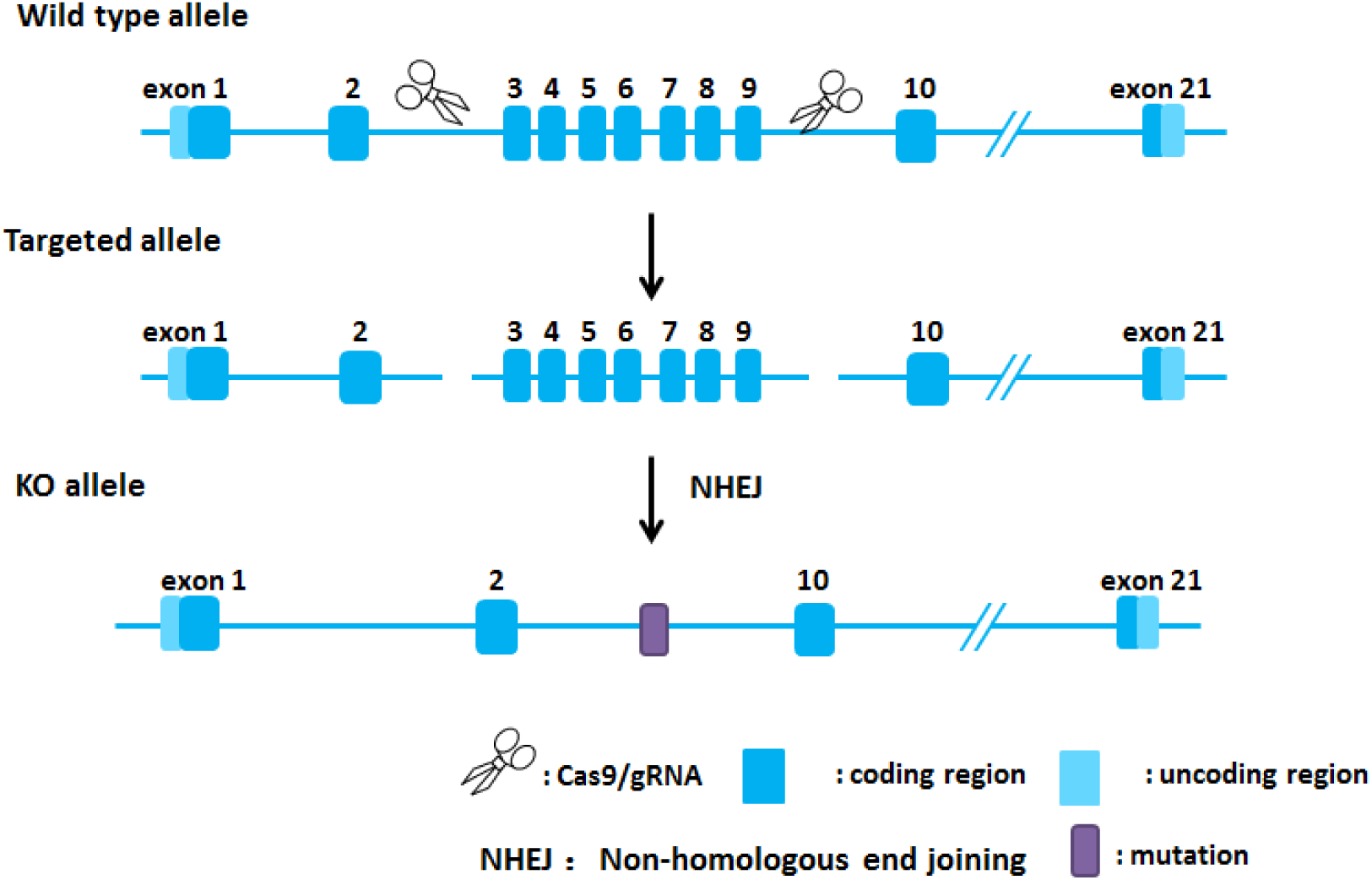
Genetic and Experimental Design ARHGEF17 knockout (KO) mice were generated using the CRISPR/Cas9-mediated gene-editing approach to introduce a frameshift mutation that disrupts the open reading frame and abolishes ARHGEF17 protein function. Briefly, Cas9 mRNA and single-guide RNA (sgRNA) targeting the Arhgef17 locus were synthesized by in vitro transcription. The Cas9 mRNA and sgRNA were co-microinjected into fertilized C57BL/6J mouse zygotes. Injected embryos were transferred into pseudopregnant female mice to generate founder (F₀) offspring. Genomic DNA from tail biopsies of F₀ mice was amplified by PCR and subjected to Sanger sequencing to confirm the presence of frameshift mutations within the Arhgef17 coding region. Five F₀ mice carrying the desired out-of-frame mutation were identified. F₀ founders were then backcrossed with wild-type C57BL/6J mice to establish germline transmission, yielding one stable heterozygous line with four F₁ offspring carrying the confirmed Arhgef17 mutation. Homozygous Arhgef17⁻/⁻ mice were subsequently obtained by intercrossing F₁ heterozygotes for experimental use.

**Figure S2.**
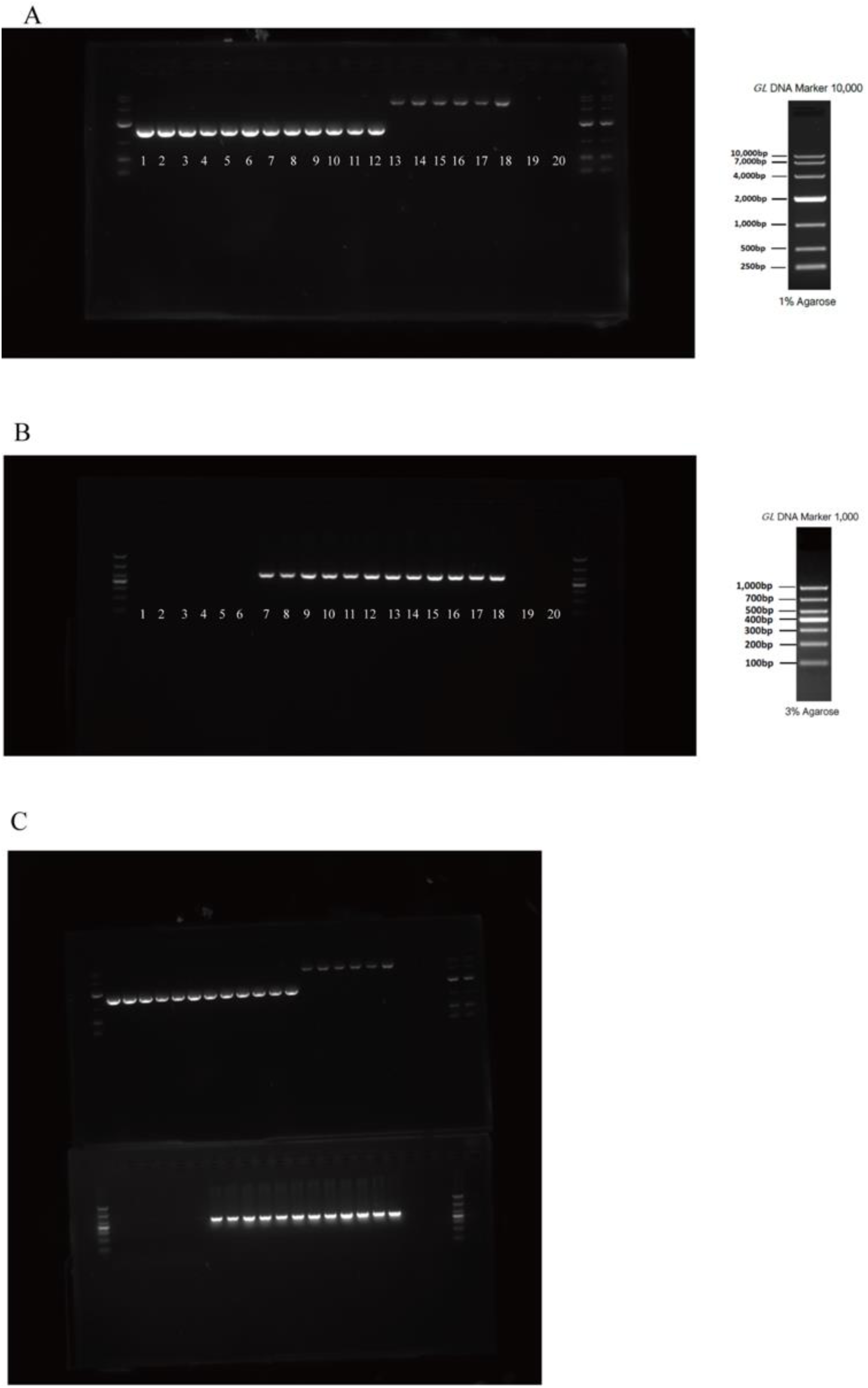
Genotype identification of ARHGEF17^-/-^ mice using agarose gel A.(P1P2) 1-12 obtain a 1.5 kb fragment, 13-18 obtain a 5.4 kb fragment.19-20 do not obtain fragment. B.(P3P4) 1-6 do not obtain a 500 bp fragment,7-18 obtain a 500 bp fragment. 19-20 do not obtain fragment. C. Merge A and B. Wild type (13-18), P1 and P2 obtain a single 5.4 kb fragment, P3 and P4 obtain a 500 bp fragment. Heterozygote (7-12), P1 and P2 obtain 5.4 kb and 1.5 kb fragments (the 5.4 kb fragment may not be obtained due to primer competition in PCR), P3 and P4 obtain a 500 bp fragment. Homozygote (1-6), P1 and P2 obtain a single 1.5 kb fragment, P3 and P4 do not obtain a 500 bp fragment. 1xTAE Buffer (19,20)

**Fig. S3.**
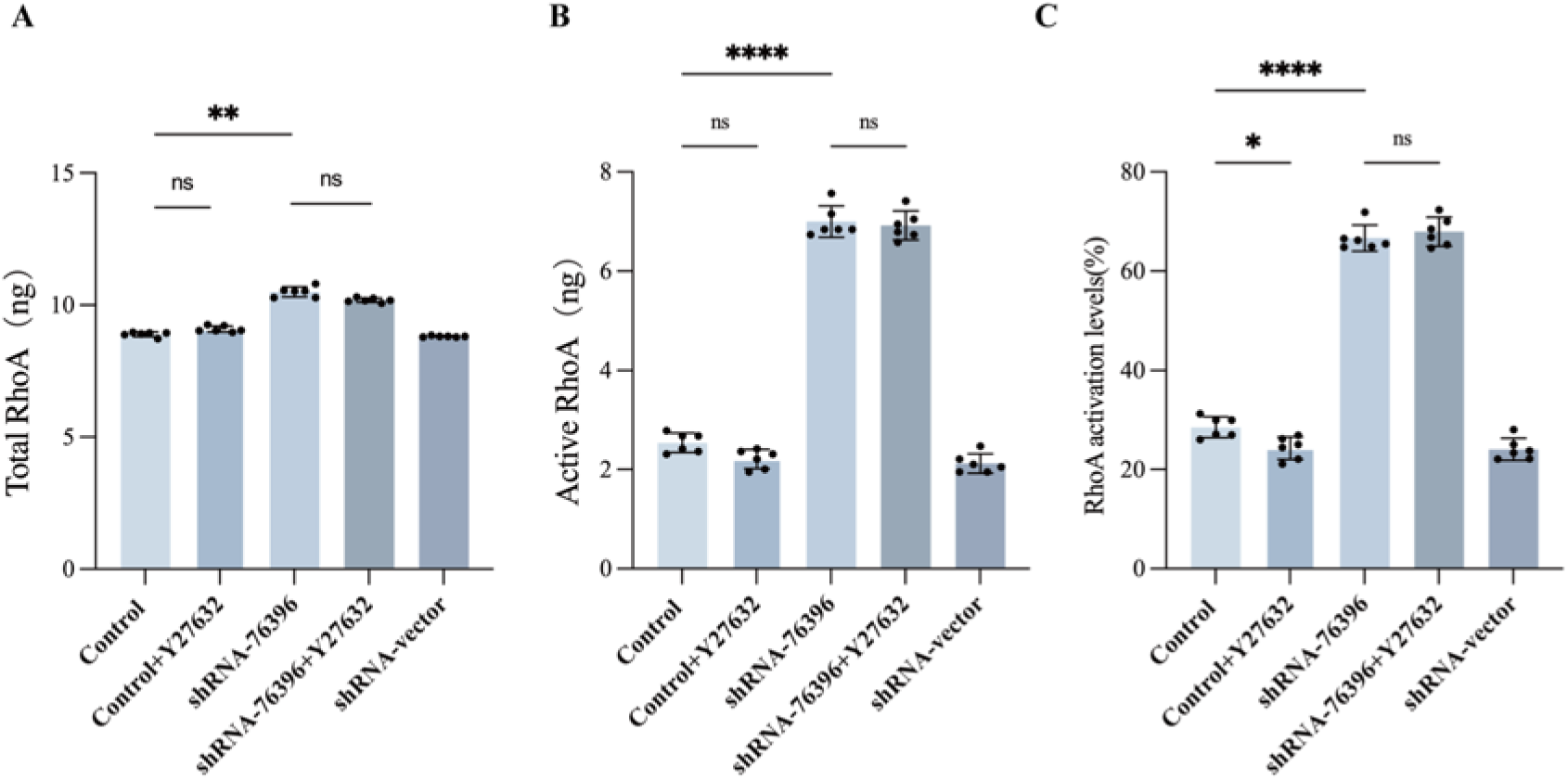
Y-27632 treatment did not reverse the shRNA-76396–induced increase in active RhoA levels or the active-to-total RhoA ratio, no significant changes were observed in total RhoA expression following Y-27632 treatment.

**Fig. S4.**
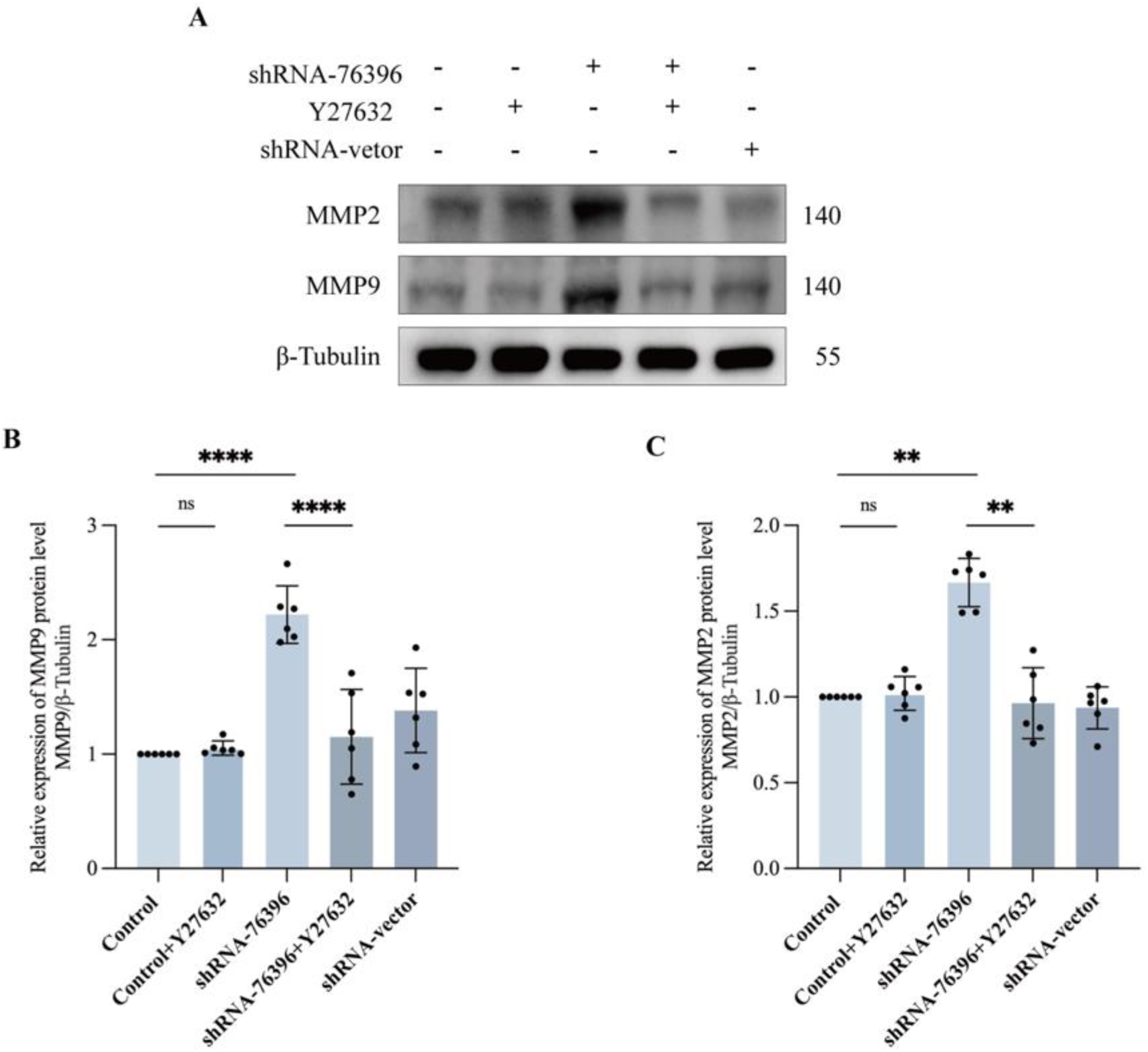
ARHGEF17 knockdown increases MMP2 and MMP9 expression in HUVECs, which is attenuated by ROCK inhibition. (A) Representative Western blot analysis showing MMP2 and MMP9 protein expression in HUVECs under different conditions: control, control + Y-27632, shRNA-76396, shRNA-76396 + Y-27632, and shRNA-vector. β-Tubulin served as the loading control. Knockdown of ARHGEF17 markedly increased MMP2 and MMP9 expression compared with control, while co-treatment with the ROCK inhibitor Y-27632 (10 μM, 24 h) partially reversed these elevations. (B–C) Quantitative densitometric analysis of MMP9 (B) and MMP2 (C) protein levels normalized to β-Tubulin. Data are presented as mean ± SD from at least three independent experiments. Statistical significance was determined by one-way ANOVA followed by Tukey’s post hoc test (**P < 0.01, ****P < 0.0001, ns = not significant).

Full unedited gel for Figure 4D

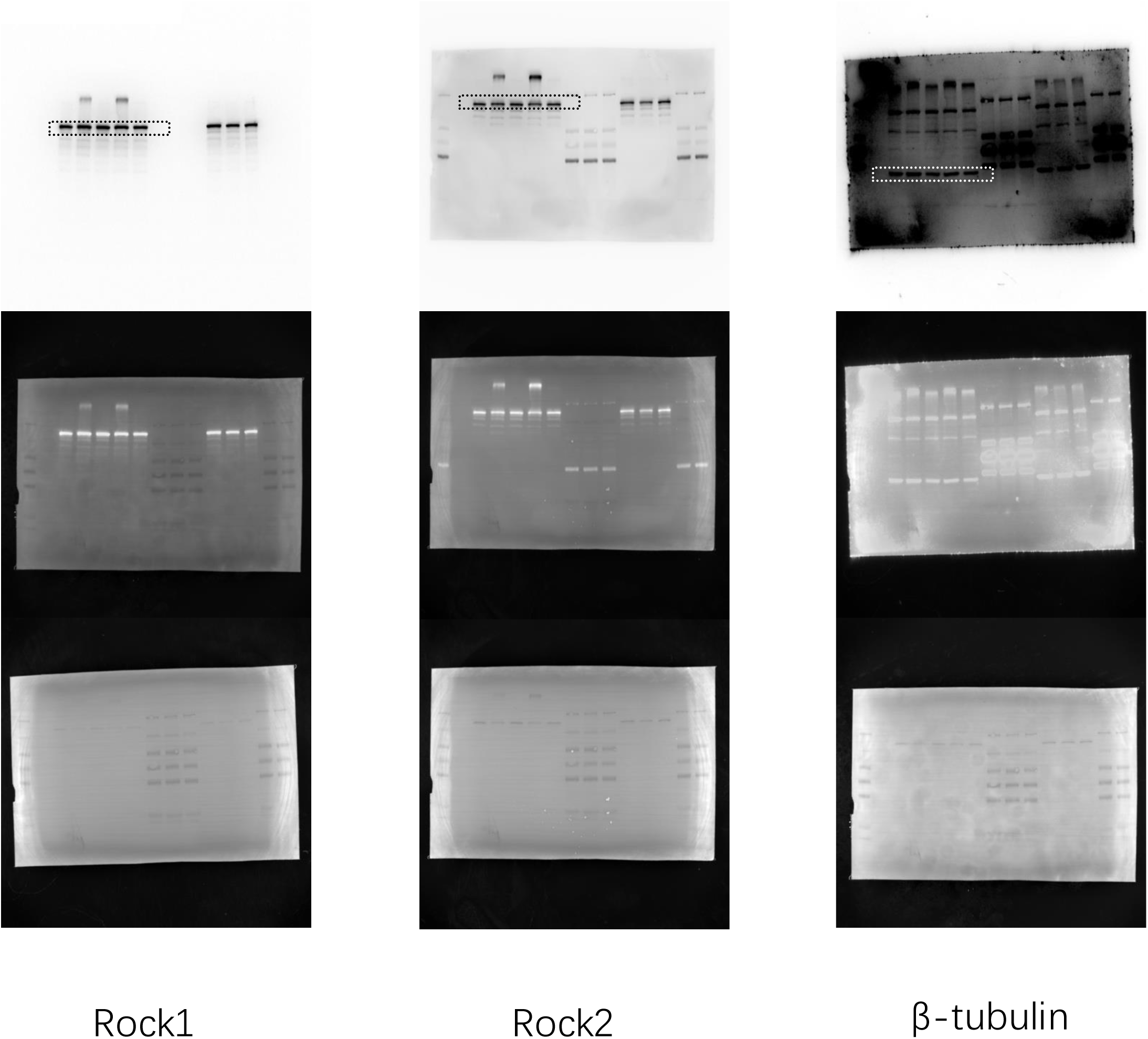

Full unedited gel for Figure 4D

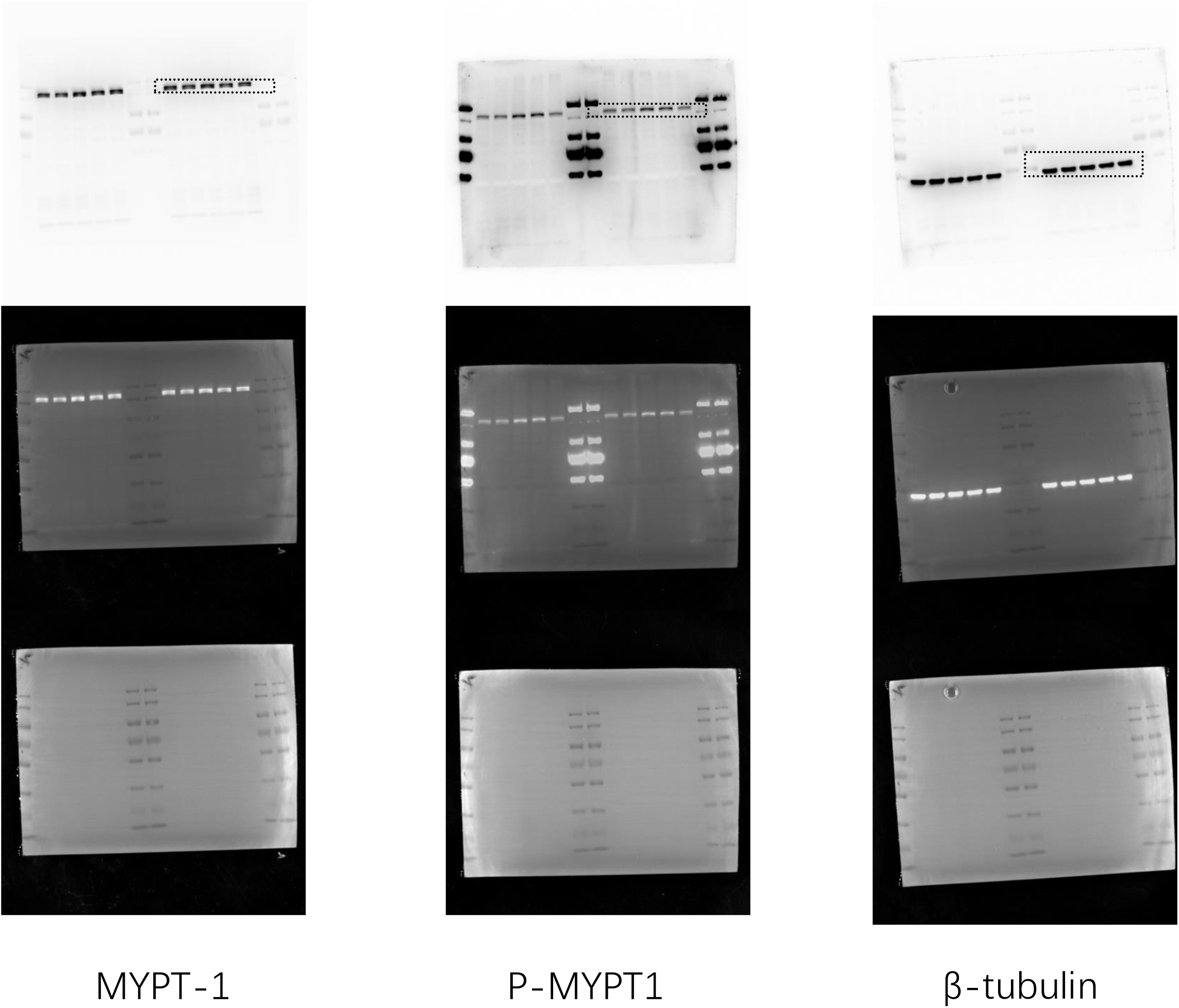

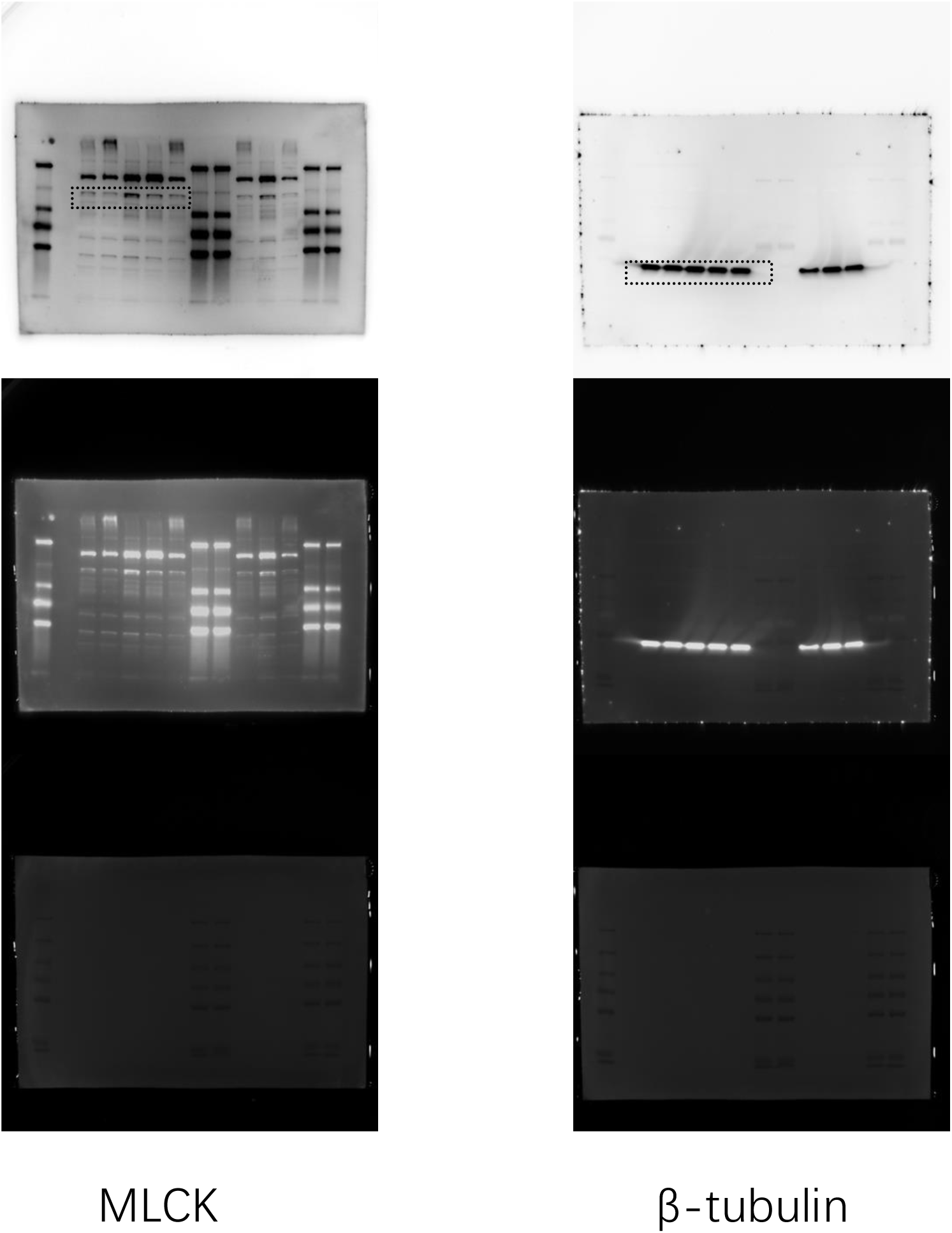

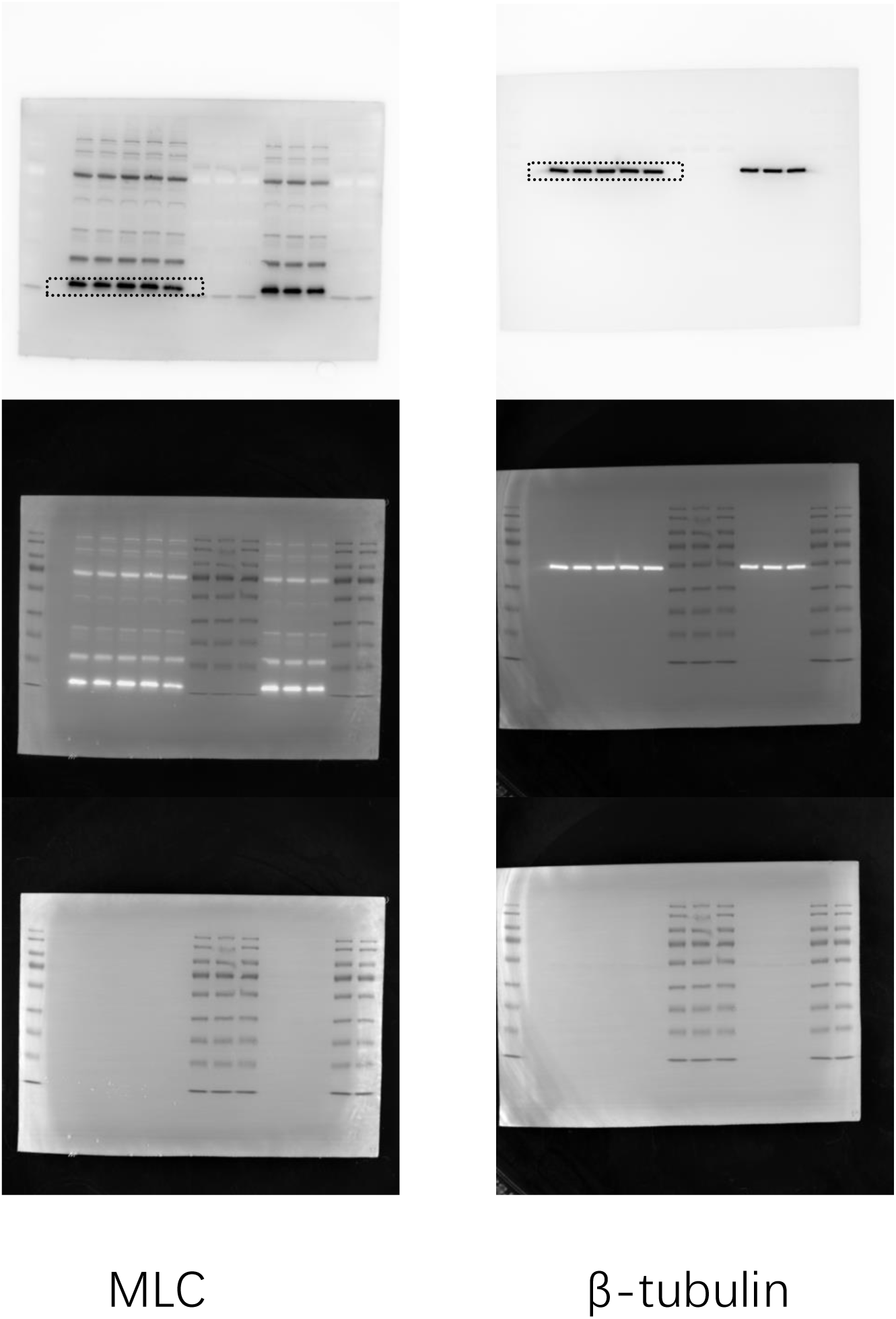

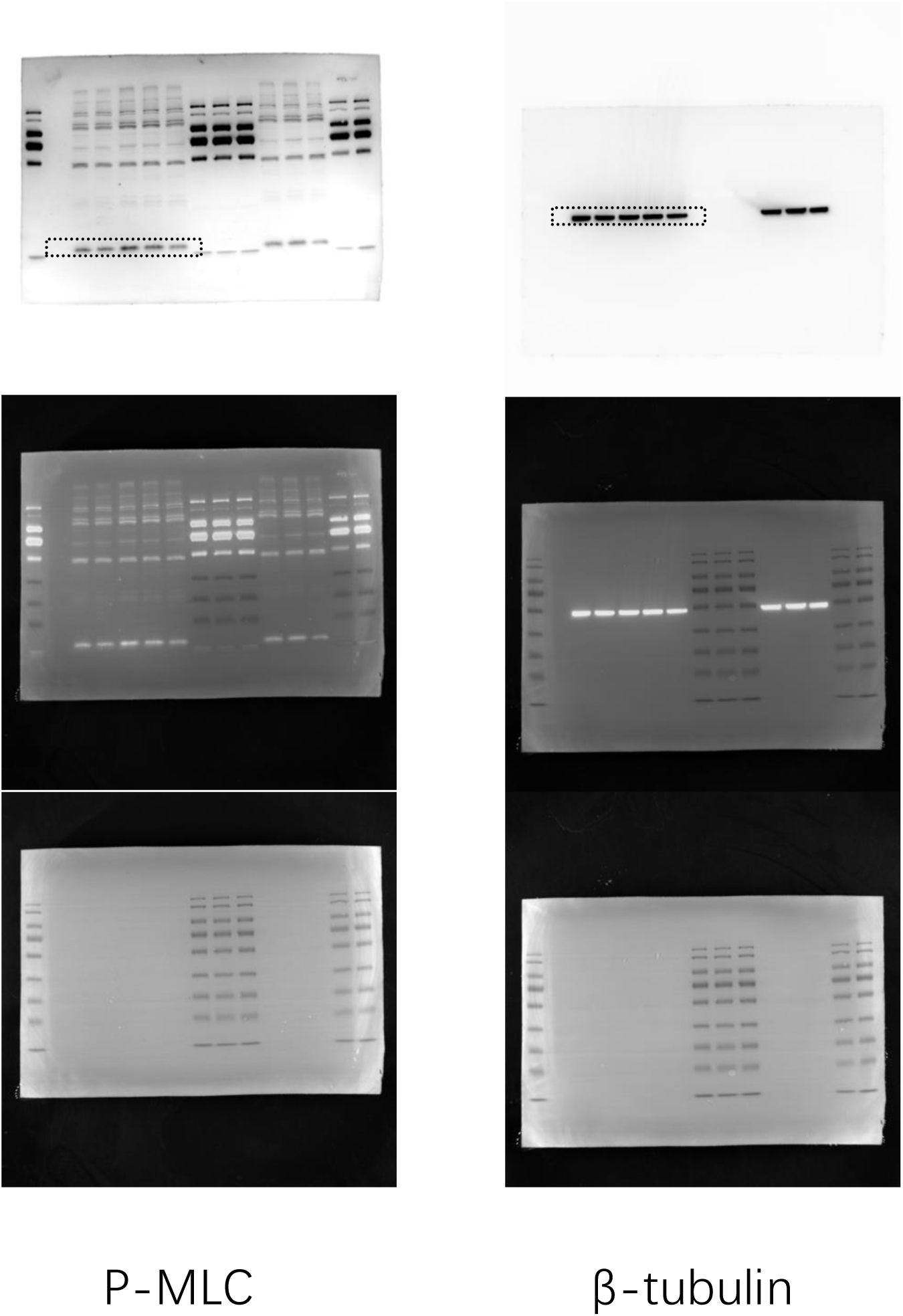

Full unedited gel for Figure 6E

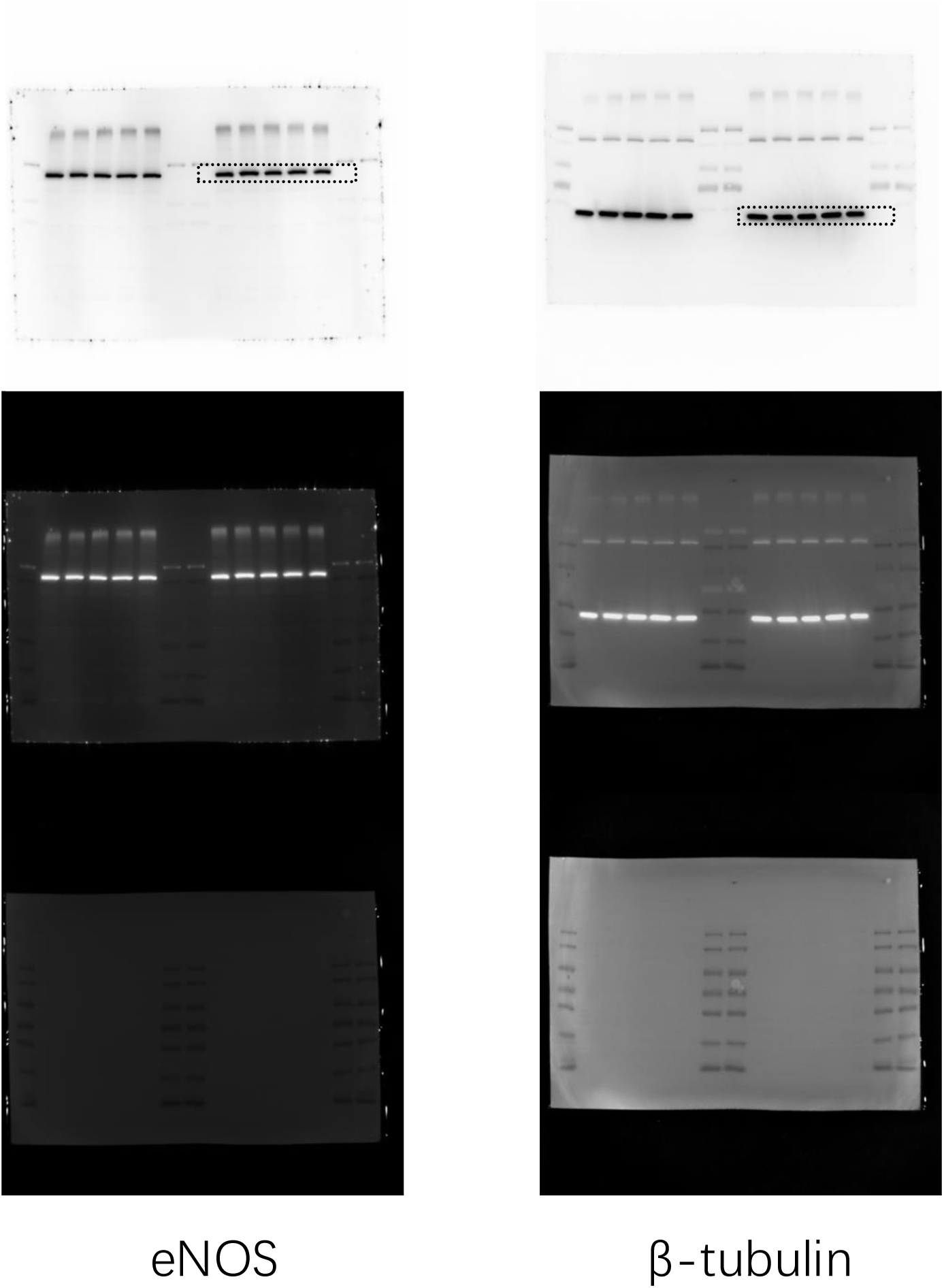

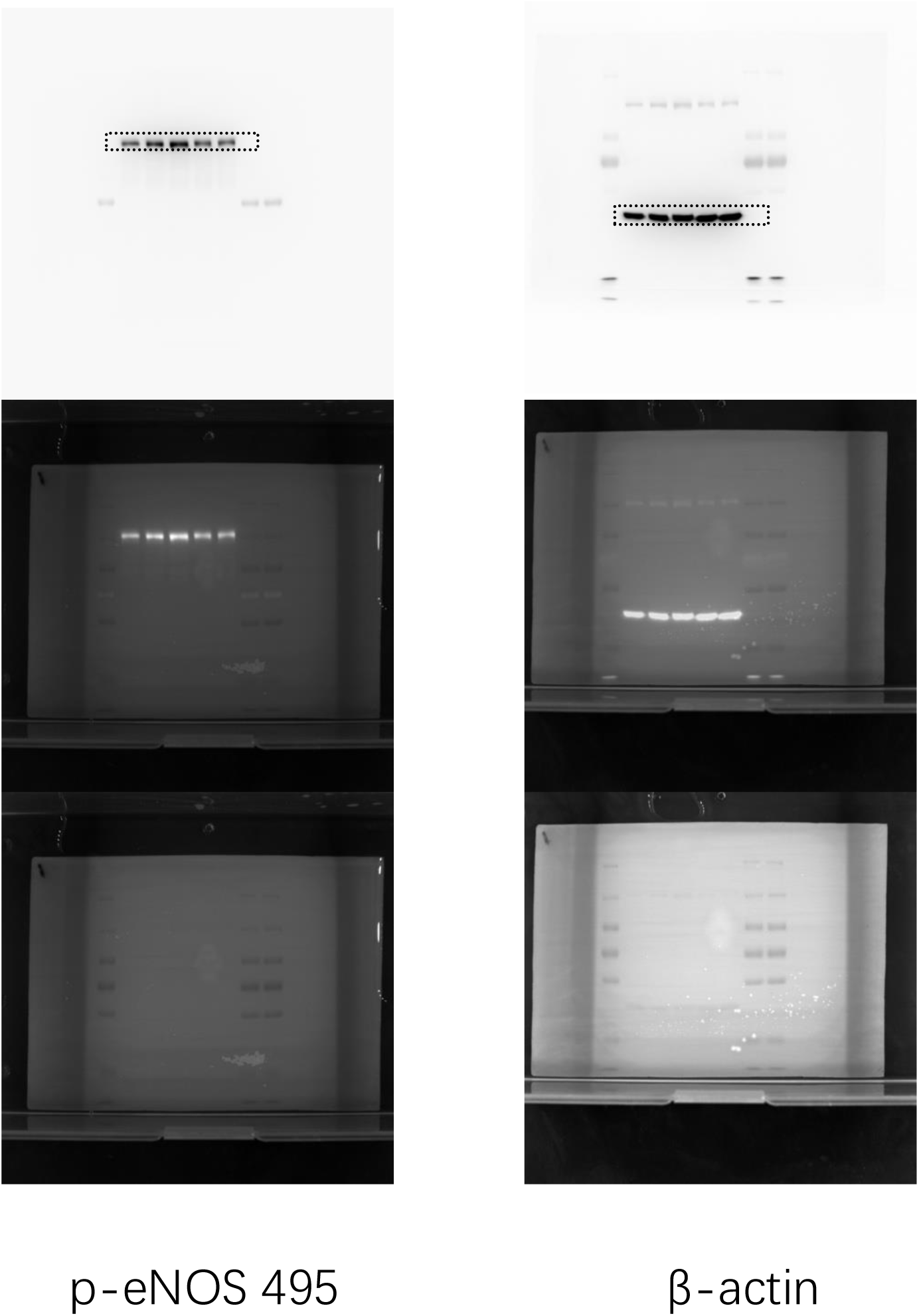

Full unedited gel for Figure 6E

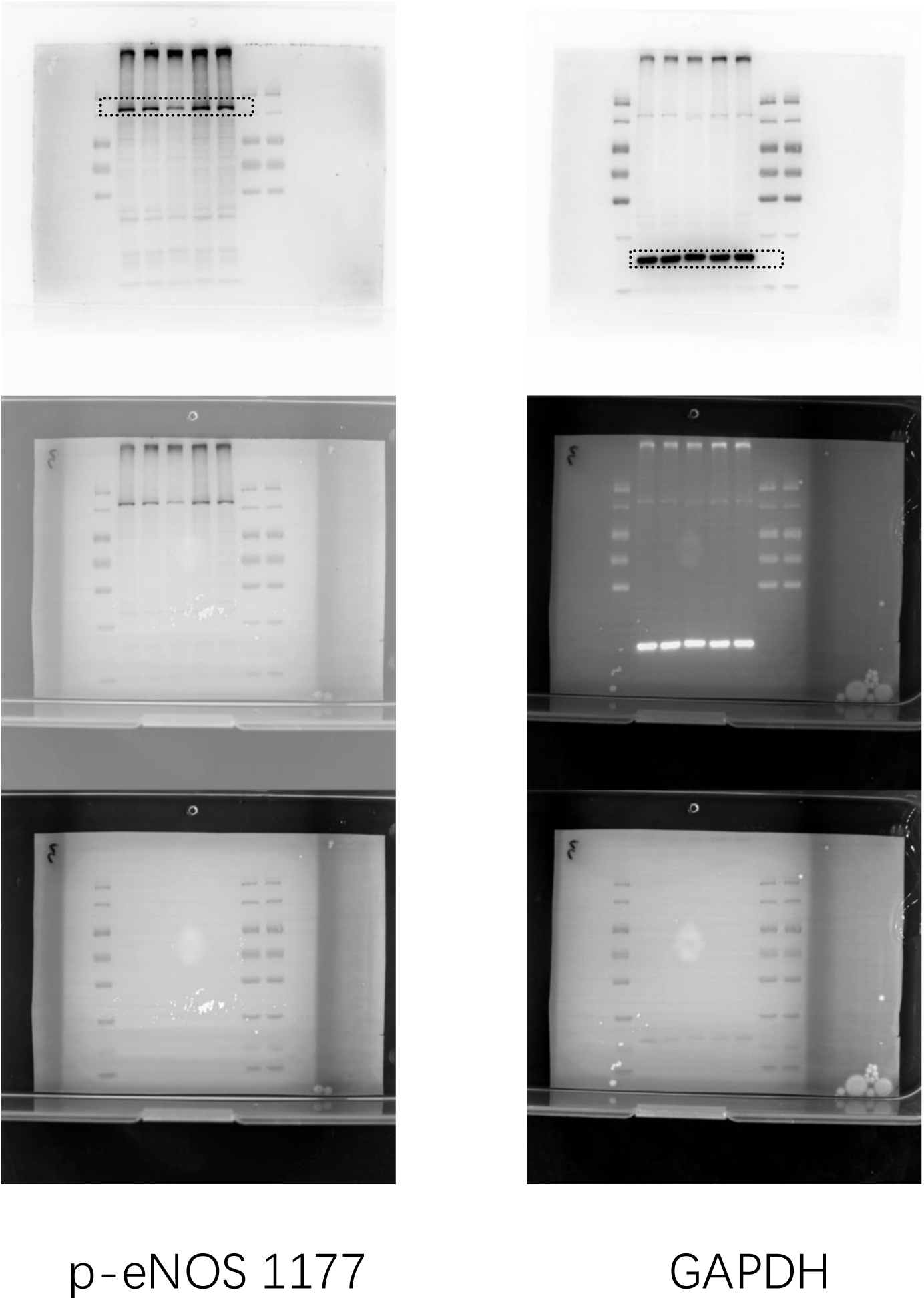

Full unedited gel for Figure S4

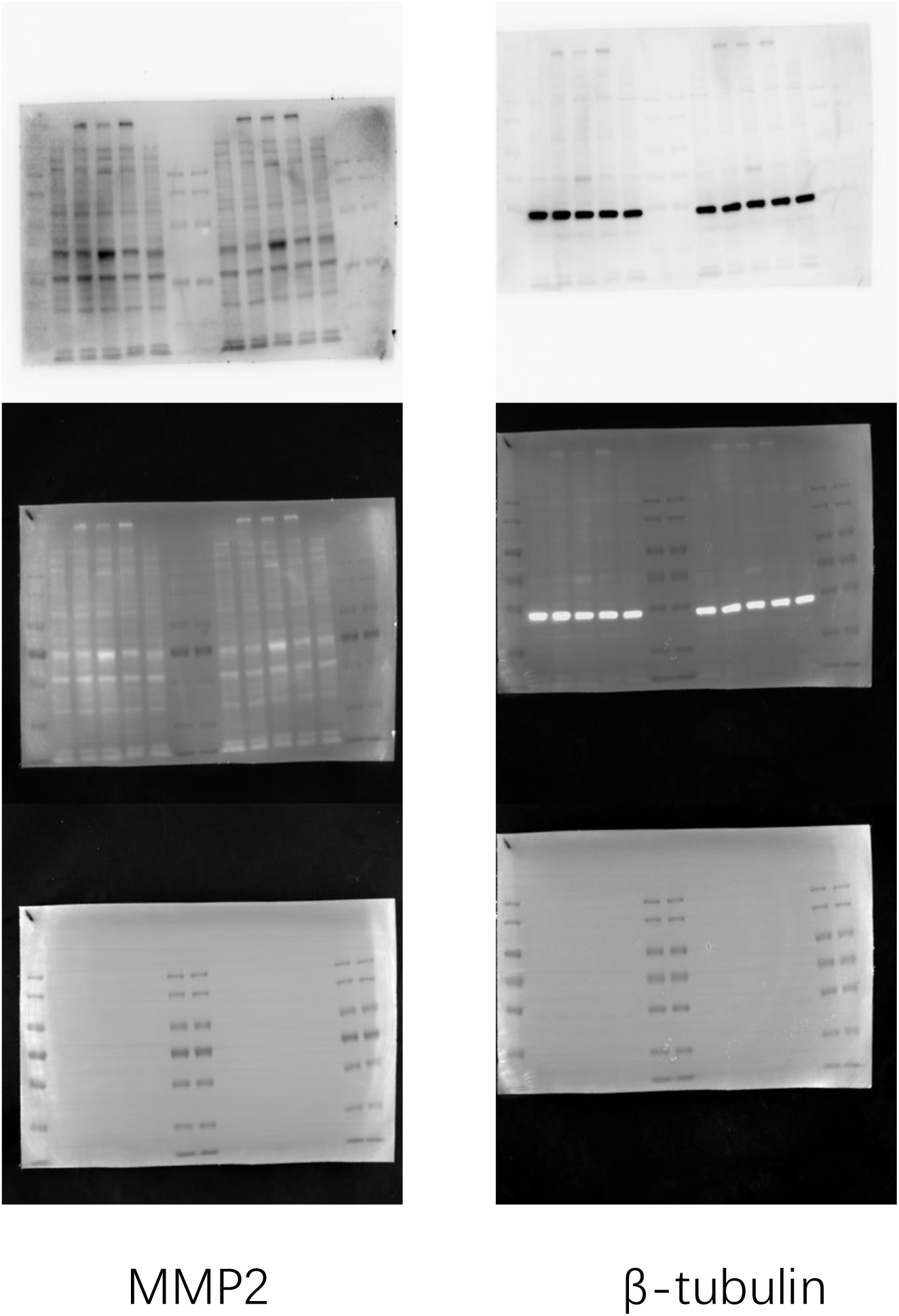

Full unedited gel for Figure S4

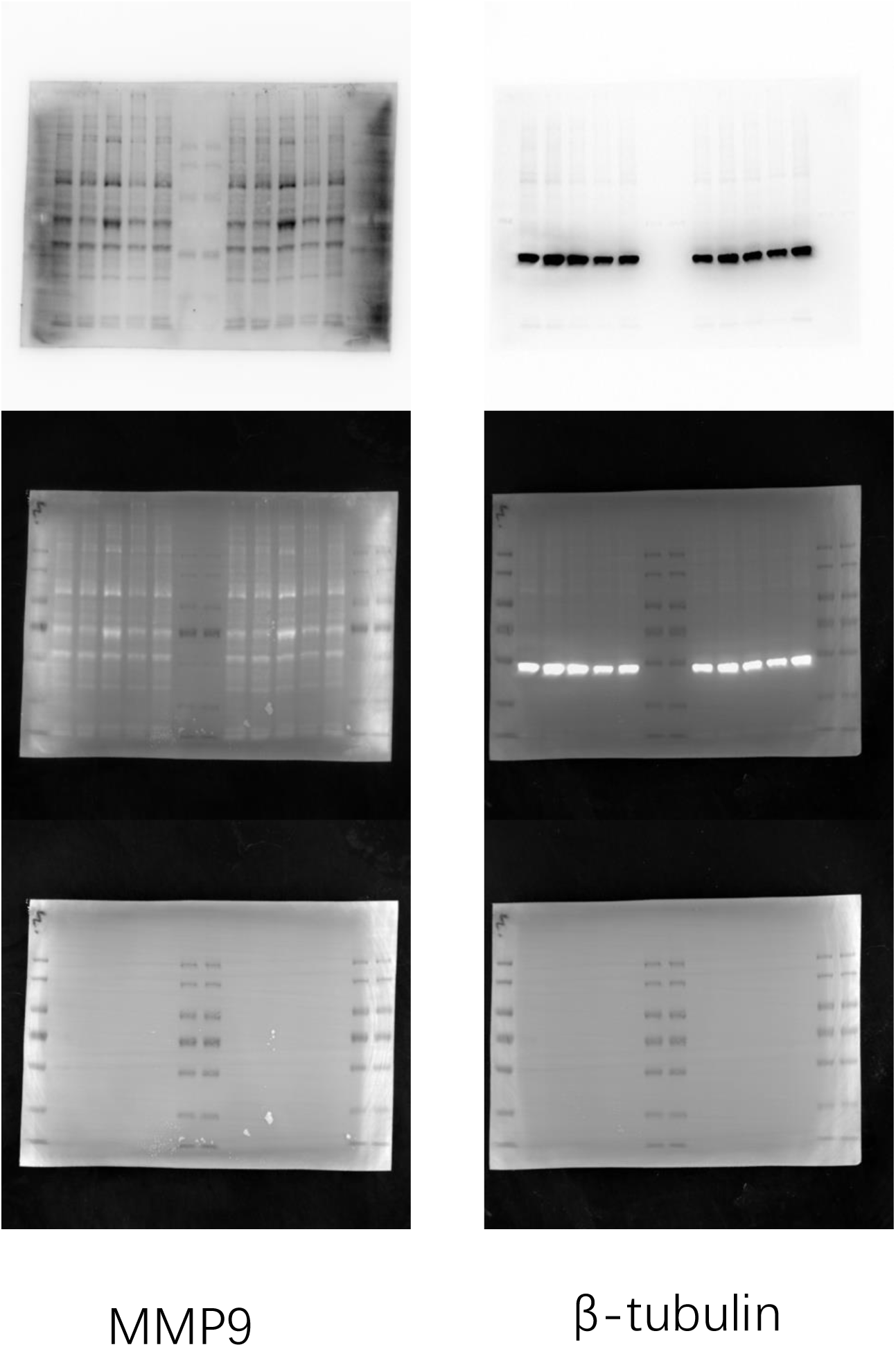

